# Cerebral microthrombi promote focal Cav-1-dependent blood-brain barrier impairment, triggering neuroinflammation and neuronal damage after traumatic brain injury

**DOI:** 10.64898/2026.09.24.754043

**Authors:** A. Wehn, M. Schifferer, J. Shrouder, A. Talukdar, M. Louessard, M. Abioui-Mourgues, V. J. Boide-Trujillo, G-M Calandra, B. Groschup, M. Reid, J. Oberhauser, U. Mamrak, M-L. Fleury, P. Marie, F. Moschogiannaki, A. Gerschmann, S. Yoshimura, S. Niedermeyer, T. Weig, N. Terpolilli, L. Dauphinot, A. Yang, A. Klymchenko, F. Ringel, D. Vivien, N. Plesnila, I. Khalin

## Abstract

Traumatic brain injury (TBI) is frequently accompanied by blood–brain barrier (BBB) dysfunction, yet the microvascular events that initiate barrier failure and secondary neural injury remain poorly understood. Using highly sensitive fluorescent nanoscale tracers, correlative light and electron microscopy, single-cell transcriptomics, and genetic manipulation of Caveolin-1 (Cav-1), we show that BBB leakage after TBI in mice occurs focally at sites of cerebral microthrombus formation. Endothelial cells adjacent to microthrombi remain structurally intact, but exhibit a distinct transcriptional program enriched in both thrombosis- and transcytosis-related genes, including Cav-1. Microthrombi promote Cav-1-dependent transcytosis and size-selective extravasation of blood-borne proteins, exposing the surrounding parenchyma to circulating factors and inducing focal microglia activation and neuronal damage *in vivo.* In human induced pluripotent stem cell-derived neurons, these proteins directly impaired neuronal structure and network function. Genetic deletion of Cav-1 markedly reduces microthrombus-associated BBB leakage, immune-cell diapedesis, microglial activation and neuronal loss, whereas endothelial re-expression of Cav-1 using AAVs restores leakage. Human TBI data reveal early dynamic coagulation dysregulation associated with adverse clinical outcome, focal cerebral vascular injury and endothelial transcriptional signatures consistent with reduced barrier integrity and increased caveolae-associated transport. Together, these findings identify cerebral microthrombi as focal sites of Cav-1-dependent BBB dysfunction and establish pathological endothelial transcytosis as a mechanistic link between post-traumatic microvascular thrombosis, blood protein extravasation and secondary neuronal injury. Targeting Cav-1-dependent transcytosis may therefore provide a strategy to limit pathological BBB permeability and secondary injury after TBI.

Graphical Abstract

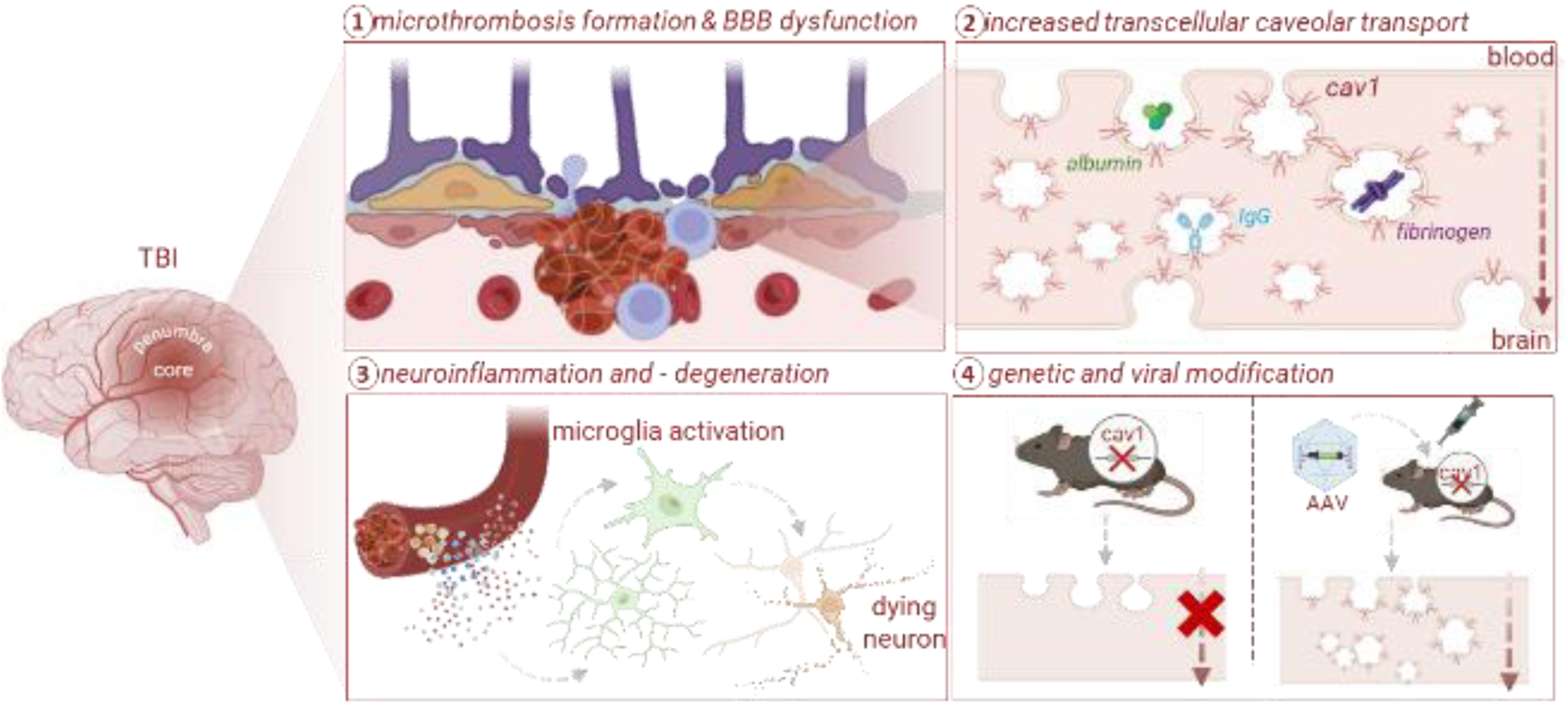

## Introduction

The blood–brain barrier (BBB) is a highly specialized interface that tightly regulates molecular and cellular exchange between blood and neural tissue thereby maintaining brain homeostasis^1^. BBB integrity relies on the coordinated function of endothelial tight junctions, regulated transcellular vesicular transport across brain endothelial cells (BECs), and perivascular support from pericytes and glial cells. Vascular injury or inflammation can compromise this barrier, leading to uncontrolled entry of circulating plasma factors that trigger neuroinflammation and neuronal damage^2, 3^. While vascular rupture is well known to induce thrombosis as a natural hemostatic response, it remains largely unexplored whether BBB dysfunction occurring in the absence of structural endothelial damage is linked to (micro)thrombi formation and whether (micro)thrombi themselves can perturb BBB function. Emerging clinical evidence indicates that microthrombi (MTi) are frequent and spatially confined events in ischemic and traumatic brain injury (TBI) and BBB disruption is also known to be independently associated with adverse brain injury outcome^4, 5^, yet their local impact on the neuro-vascular unit remains unclear^6, 7^. Here, we identify a close spatial association between MTi and focal BBB dysfunction and show that MTi-related local BBB impairment occurs through Caveolin-1 (Cav-1)–dependent transcytosis, leading to size-dependent extravasation (EV) of blood-borne proteins. Using complementary genetic, ultrastructural and functional approaches, we further examine how this focal vascular response influences the surrounding neurovascular environment and assess its relevance in human TBI.

## Results

### Association of microthrombi and focal BBB dysfunction

BBB dysfunction across neurological disorders, including TBI, is typically described as a diffuse process, largely based on the parenchymal distribution of different molecular tracers^8^. We found that the low–molecular weight tracer Cascade Blue dextran (10 kDa) shows widespread distribution throughout the traumatic contusion 2 h after controlled cortical impact (CCI; Fig. 1a). In contrast, systemically co-injected 30 nm fluorescent lipid nanodroplets (LNDs) revealed discrete sites of BBB permeability confined to specific vascular segments (Fig. 1a-b). These focal LND signals are concentrated within the traumatic penumbra, as evidenced by lectin staining (Fig. 1b) and predominantly associated with elongated, rod-like intravascular structures, indicative of microvascular occlusions. To resolve the cellular architecture of these leakage sites, we analyzed our previously published, open-source correlative light and electron microscopy (CLEM) dataset of 2 h post-TBI mouse brain (Schifferer et al, 2024^9^ from Kislinger et al., 2024^10^) and relocated a focal extravasation site from confocal microscopy to scanning electron microscopy (SEM). High-resolution SEM revealed that LNDs extravasation occurred across BECs with preserved integrity directly next to fibrin, platelets, and immune cells, indicating a transcellular transport route in the absence of tight junctions (TJs) disruption or structural endothelial damage (Fig. 1c). These observations suggest that early BBB permeability after TBI occurs focally at sites of (micro)thrombus formation (Fig. 1d).

**Fig. 1.**
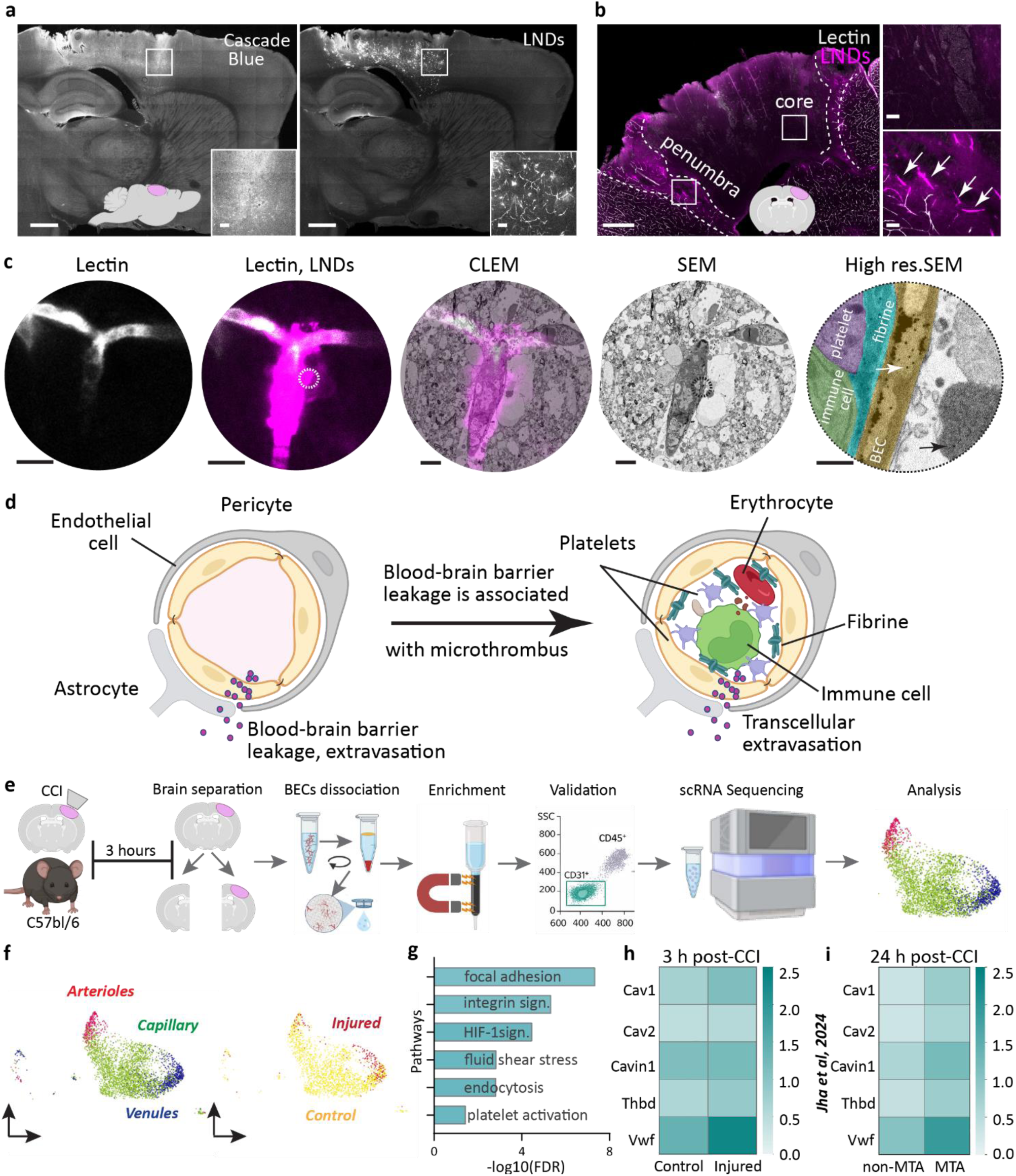
Blood-brain barrier leakage is associated with formation of microthrombus. **a,** Representative sagittal confocal microscopy image of a C57/Bl6 mouse brain after controlled cortical impact (CCI), showing different pattern of the blood-brain barrier (BBB) leakage between Cascade Blue and highly-fluorescent 30 nm lipid-nanodroplets (LNDs). Scale bar: 1mm; insert: 100 µm. **b,** Representative coronal confocal image of a mouse brain after CCI, showing LNDs accumulated in the penumbra both inside and outside (white arrows) vessels (lectin), visualizing occlusions and areas of BBB leakage. Scale bar: 500 µm; insert: 50 µm. **c,** Correlative light and electron microscopy (CLEM). Region of interest containing extravasation of LNDs from brain vessels was relocated and imaged by scanning electron microscopy (SEM). High resolution SEM of ROI in SEM overview image (dashed circle) showing extravasation reveals the presence of fibrin (cyan), platelet (magenta) and immune cell (green) at the site of extravasation and transcytosis of nanoparticles (NPs) via visually non-damaged brain endothelial cell (BEC). Black arrow - NPs in the brain; white arrow - NPs travelling across BEC. Scale bars (from left to right): 10 µm; 10 µm; 5 µm; 5 µm; 500 nm. Original raw images are from Schifferer et al^9^. **d,** Schematic illustrating the conventional paradigm shift towards the association between BBB leakage and microthrombi. **e,** Schematic representation of the experimental workflow for BEC scRNA sequencing in mice 3 hours after CCI. **f,** UMAP clustering of arterioles, venules, capillaries, and identification of injury-associated endothelial cells using Immuno-MRI (Ext. data Fig.1 a). **g,** KEGG pathway enrichment analysis of differentially expressed genes (DEGs). **h–i,** Color-coded grid maps illustrating increased expression of individual genes involved in transcytosis and microthrombus formation at 3 hours post-CCI in the original dataset **(h)** and at 24 hours post-CCI in a bulk RNA-seq dataset reported independently by Jha M. et al.^11^ **(i).**

To test whether this phenotype is accompanied by corresponding transcriptional changes, we performed single-cell RNA sequencing of BECs in mice at 3 h post-CCI, matching the timing of this early event (Fig. 1e). Uniform Manifold Approximation and Projection (UMAP) projection identified arteriolar, venular and capillary endothelial cells (Fig. 1f), consistent with established zonation markers. Given the focal nature of the CCI model (3 mm diameter, 1 mm depth), which produces a relatively restricted lesion and penumbra (Fig. 1a-b), only a small proportion of BECs are involved in microthrombus formation and BBB leakage. Therefore, direct ipsilateral versus contralateral comparisons were not performed. Rather, activated or “Injured” endothelial cells were defined within the ipsilateral hemisphere based on *Icam1* and *Vcam1* expression, while contralateral BECs served as controls (Fig. 1f). To confirm that these cells are associated with injured tissue, longitudinal molecular MRI was performed to reveal an early increase in VCAM-1 expression in perilesional vasculature, implicating rapid endothelial activation after injury (Extended Data Fig. 1 a). KEGG pathway enrichment analysis of differentially expressed genes identified that injured BECs were associated with endothelial mechanotransduction (Focal adhesion, Integrin signaling, Fluid shear stress), hypoxic stress (HIF-1 signaling), vesicular trafficking (Endocytosis), and platelet interactions (Platelet activation), suggesting activation of endothelial programs associated with microthrombus and BBB dysfunction (Fig. 1g, Extended Data Fig. 1 b). Analysis of individual functional gene programs also revealed increased expression of transcytosis-and thrombosis-related genes at 3 h post-injury, including *Cav1*, *Cav2*, *Cavin1, Thbd* and *Vwf* (Fig. 1h-i, Extended Data Fig. 1 c).

This coordinated activation suggests that microthrombus formation and gene expression in BECs are closely associated with increased expression of genes promoting transcytosis across the BBB. To examine how this response evolves over time and validate independently, we re-analysed an open-source bulk RNA-seq dataset of mouse brains 24 h after CCI^11^. Subclustering of endothelial cells (Extended Data Fig. 2 a-b) revealed marked induction of *Spp1* expression at 24 h post-TBI (Extended Data Fig. 2 c-d), identifying a distinct transcriptional subset of injury-associated BECs. This population was further characterized by elevated expression of *Icam1* and *Ccl2* (Extended Data Fig. 2 e). Within this subset, a small subcluster (Extended Data Fig. 2 f) also exhibited pronounced upregulation of caveolar transcytosis genes (*Cav1, Cav2, Cavin1*) together with microthrombus-associated (MTA) genes (*Vwf, Thbd*) compared to other injured BECs (Fig. 1h). We termed these cells MTA BECs, representing a distinct endothelial state associated with MTi-related BBB leakage. Further analysis of this subcluster revealed co-expression of gap junction proteins *Gja1* and *Gjc1*, implicating post-capillary venules as particularly vulnerable sites for MTi formation and BBB disruption (Extended Data Fig. 2 g). In addition, MTA BECs showed coordinated upregulation of thrombosis-related genes (*Vwf, Thbd, Vwa1, Vtn*; Extended Data Fig. 1h), alongside signatures of vascular stress, inflammation, a pericyte-like transition (*Pdgfrb, Rgs5*)^12^ and BBB dysfunction. Together, these findings suggest that early after TBI, MTi formation is associated with a distinct endothelial transcriptional landscape characterized by enhanced transcytosis, pro-thrombotic signaling and a further shift from a BBB-specialized phenotype toward a transitional, pericyte-like state with impaired barrier function^13^. Notably, these molecular changes occurred despite preserved endothelial morphology, consistent with imaging data demonstrating BBB leakage exclusively in morphologically intact vessels occluded by MTi (Fig. 1a–c).

### Morphological and functional characteristics of microthrombi-mediated BBB dysfunction

In our recent studies^10, 14^, we identified MTi associated with focal BBB leakage in the traumatic penumbra. The CCI thus provides a well-defined spatial and temporal *in vivo* model to study MTi formation and BBB dysfunction alongside TBI. Systemically injected 30 nm LNDs 1 h after CCI enabled simultaneous visualization of these processes (Fig. 2a) through intravascular accumulation (MTi) and extravasation (BBB dysfunction). As previously described, our LNDs exhibit a highly bright fluorescence due to excitation energy transfer between loaded fluorophores^14^. Such properties enable us to detect even small vascular leakage sites. To preserve the high fluorescence of the LNDs during standard histological processing and maximize sensitivity, we established a correlative confocal microscopy (CCM) workflow (Fig. 2a) in which the same brain samples were imaged twice: prior to and after preparing the samples for immunohistochemistry (IHC). This approach retained the native LND signal and enabled their accurate co-localization with vascular and cellular markers (Fig. 2a).

**Fig. 2.**
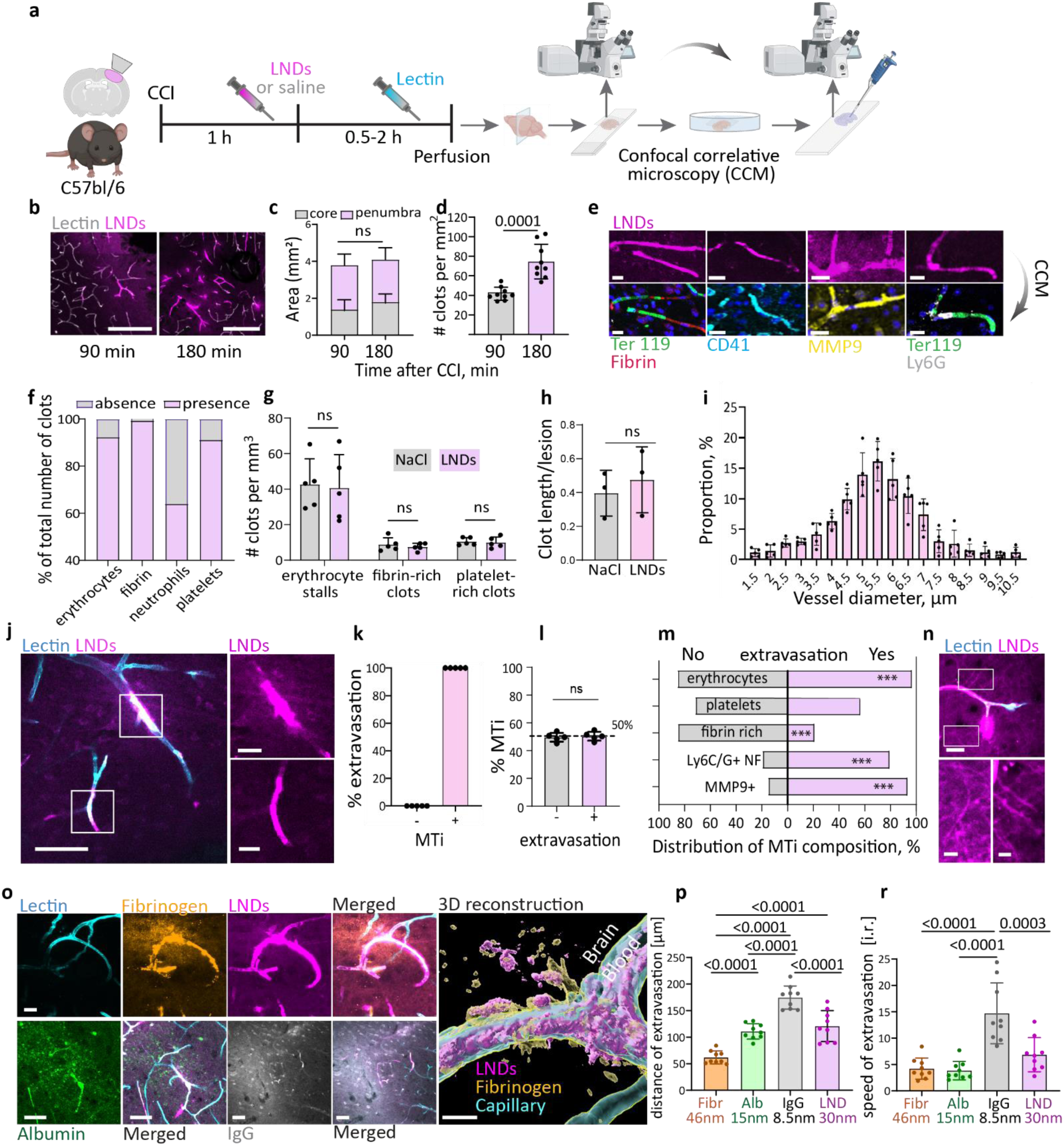
Structural and functional particularities of microthrombi-mediated BBB dysfunction. **a,** Experimental pipeline: 1 hour after CCI, mice were injected i.v. with 30 nm LNDs or saline, followed by lectin injection, perfusion and tissue collection. **b,** Confocal microscopy images showing the distribution of LND-labelled microthrombi (MTi) in penumbra 90-or 180-min post-CCI. Scale bar: 200 µm. **c**, Analysis of penumbra and lesion core areas in different time-points post-TBI. **d,** Analysis of MTi appearance post-CCI. Mean ± SD; n=5 mice, unpaired t-test. **e,** Confocal correlative microscopy (CCM) reveals a variety of LNDs-labelled MTi compositions. Scale bar: 10 µm. **f,** Analysis of proportional distribution of MTi components. n=161/340/360/427 clots. **g-h,** Analysis of influence of LNDs on MTi formation and size. Mean ± SD; n=5/3 mice, unpaired t-test. **i,** Size distribution of the vessels with MTi. Mean ± SD of proportion of vessel diameter with detected MTi. n = 62 clots (6 mice). **j,** Images showing LND-labelled MTi with (upper) and without (lower) extravasation. Scale bar: 50 µm; zoomed 10 µm. **k,** Analysis of the proportion of extravasation sites relative to MTi. **l,** Analysis of proportional number of MTi with and without extravasation. Mean ± SD; n=5 mice, unpaired t-test. **m,** Regression analysis of relation between MTi composition and blood-brain barrier dysfunction. n=161/340/360/427 clots, Pearson correlation analysis, ***<0.0001. **n,** Images showing accumulation of extravasated LNDs inside neuronal fibers. Scale bar: upper 20 µm; lower 5 µm. **o,** Images of accumulation and extravasation of fluorescently labelled blood-borne components (fibrinogen, albumin, IgG) and LNDs at MTi. 3D reconstruction showing LNDs and fibrinogen extravasation. Scale bars: fibrinogen – 20 µm; albumin, IgG – 50 µm, 3D – 5 µm; **p-r,** Analysis of the distance (**o**) and speed (**p**) of extravasation for fibrinogen (orange), albumin (green), IgG (grey) and LNDs (magenta) shows size-selectivity. n=5 mice per group, One-way ANOVA.

Using this pipeline, we first identified two spatially distinct regions after CCI: a penumbral zone containing both lectin-labelled vasculature and focal LND signals, and a lesion core largely devoid of lectin staining. Both the penumbra and the lesion core showed a trend of expansion between 1.5 h and 3 h following TBI (Fig. 2b-c). In contrast the number of LND-labelled MTi significantly increased within the penumbra, reaching nearly two-fold higher levels at 3 h post-CCI (Fig. 2d), suggesting that this process may contribute to secondary penumbral damage. Morphologically, MTi exhibited heterogeneous cellular compositions, with variable amounts of platelets, fibrin, erythrocytes, and leukocytes (Fig. 2e-f). Importantly, administration of LNDs did not affect the number or size of MTi (Fig. 2g-h), validating LNDs as coagulation-inert imaging probe.

Further, analysis of vessel diameter showed that MTi occurred predominantly in capillaries (3-4 µm) and post-capillary venules (8-9 µm; Fig. 2i)^15^, consistent with our transcriptomic analysis (Fig. 1f; Extended Data Fig. 2 g). Extravasation of LNDs was observed exclusively in vessels containing MTi and was never detected in MTi-negative vessels (Fig. 2k). However, only approximately half of MTi identified were associated with measurable extravasation (Fig. 2l). These data indicate that BBB leakage is not an obligate consequence of microthrombus formation. Instead, our recent findings lead us to speculate that prolonged retention of MTi within microvessels may result in progressive, time-dependent BBB activation and extravasation, as suggested by pseudotime clot reconstruction in our previous study^16^. To identify features of MTi associated with leakage, we examined the cellular and molecular composition of MTi 90 min post-TBI. Platelets and erythrocytes were present in both leaking and non-leaking MTi, whereas dense fibrin content was negatively associated with LND extravasation. By contrast, the presence of immune cells and/or MMP9 expression markedly increased the likelihood of leakage, as revealed by correlation analysis (Fig. 2m).

Collectively, these findings indicate that MTi composition shapes the propensity for BBB leakage, highlighting a functional interplay between blood-derived components and endothelial barrier responses. Moreover, the trajectory of EV events suggests that LNDs are initially accumulate within MTi and then subsequently detected crossing the adjacent endothelium and later entering the surrounding parenchyma. Once extravasated, LNDs are not confined to the perivascular space but spread into the surrounding parenchyma, extending into fine, fiber-like structures displaying bouton- and spine-like morphologies, characteristic of neuronal dendrites and axons (Fig. 2n), suggesting that neurons are directly exposed to extravasated blood-borne material.

To determine whether endogenous blood components exhibit a similar distribution as LNDs, we examined post-CCI parenchymal localization of plasma-derived proteins. Regions of LND leakage overlapped with EV areas of systemically injected fluorescently labelled blood-borne proteins, including fibrinogen, albumin, and IgG (Fig. 2o). Similar to LNDs, these proteins accumulate at MTi sites before extravasating into the parenchyma, suggesting that occlusions act as focal accumulation hotspots that additionally facilitate transvascular entry of blood-borne proteins into the brain parenchyma. EV exhibited clear size-dependent kinetics: the smaller molecules, e.g. IgG, extravasate more rapidly and over greater distances as compared to larger molecules, e.g. fibrinogen (Fig. 2o–p). Notably, such focal EV hotspots may locally expose neuronal networks to high concentrations of inflammatory mediators and thereby contribute to neuronal dysfunction. Importantly, in our previous studies we observed enhanced vesicular structures within BECs surrounding persisting and leaking MTi, consistent with vesicle-mediated transport processes^10, 16^. This raises the possibility that microthrombi-associated BBB dysfunction is not solely driven by passive barrier breakdown, but instead involves active, transcellular mechanisms. Additionally, we previously observed elevated Cav-1 expression near microvascular occlusions in models of stroke and traumatic injury,^14, 16^ supported by our current data on scRNA transcriptomics of BECs (Fig. 1 g-i). This prompted us to investigate the role of caveolae-mediated transcytosis and Caveolin-1 (Cav-1) in microthrombi-driven BBB dysfunction.

### Cav-1 drives microthrombus-associated BBB dysfunction

Cav-1 is a key regulator of endothelial caveolae-mediated transcytosis and has been implicated in BBB permeability under several pathological conditions, including aging^17^, neuroinflammation^18^, and acute brain injury^19^. We found that Cav-1 expression increased markedly in the ipsilateral cortex following CCI and was enriched in endothelial cells associated with microthrombi at transcriptomic level (Fig. 3a–c; Fig. 1g-h). To directly assess the functional role of Cav-1 in MT-associated BBB dysfunction, we compared Cav-1 wt and ko mice injected with 30-nm LNDs or the blood-borne protein albumin at acute (2 h) and sub-acute (48 h) stages after CCI (Fig. 3d). We first confirmed that Cav-1 ko mice (Ext. data Fig. 3a) exhibit a significantly reduced Cav-1 expression in brain endothelial cells compared to Cav-1 wt controls (Ext. data Fig. 3 a-b). Furthermore, we demonstrated that systemic Cav-1 deletion did not affect the circulation kinetics of LNDs during the time window used for MTi and BBB leakage labelling (Extended Data Fig. 3c–d). In our experimental setup (Fig. 3d), we confirmed that the total fluorescent intensity of LNDs from injured brain slices (accumulated inside MTi and extravasated) was comparable between Cav-1 wt and Cav-1 ko mice at both time points (Extended Data Fig. 3e-h). This indicates that similar amounts of LNDs are retained in the brain of both experimental groups after TBI, which allows a valid comparison of MTi formation and LNDs extravasation across genotypes. Using this approach, we observed that MTi formation was prominent early (2h) following TBI and declined over two days in both wt and ko animal groups (Fig. 3e). Cav-1 deficiency was associated with fewer MTi at 2 h post-injury, an effect that was no longer evident at 48 h (Fig. 3e). Strikingly, the proportion of MTi associated with leaky BBB was reduced by more than half in Cav-1 ko mice and remained significant at both time points (Fig. 3f), identifying Cav-1 as a key mediator of transcellular MTi-BBB dysfunction. Analysis of singular microclots revealed a clear re-distribution of LNDs within MTi depending on Cav-1 expression (Fig. 3g–h). In wild-type animals, approximately 80% of LNDs extravasated into the perivascular parenchyma, with only ∼20% remaining within the clot. Genetic deletion of Cav-1 reduced both albumin and LND extravasation at MTi, indicating that pathological activation of Cav-1 not only amplifies endothelial transcytosis but also broadens the range of transported molecules (Fig. 3g–h). Thus, in the absence of Cav-1, a larger fraction of blood-borne components remained confined within the vasculature, thereby limiting exposure of the surrounding brain parenchyma to blood-borne components. Interestingly, at 48 h post-CCI we no longer observe differences in the extravasation of either albumin or LNDs between Cav-1 wt and Cav-1 ko mice, indicating that Cav-1–dependent transcytosis is no longer the predominant pathway in this later stage (Fig. 3g–h). Instead, BBB leakage at 48 h is likely dominated by paracellular mechanisms associated with TJ degradation, which was observed in both genotypes (Extended data Fig.3 i-j). Additionally, at 48h after injury, extravasated LNDs and albumin were internalized by CD45⁺ immune cells in the peri-lesional area (Extended Data Fig. 3k-l), suggesting that blood-borne components no longer directly affect neurons but get neutralized by infiltrating peripheral immune cells. Cumulatively, these data suggest pathological upregulation of Cav-1 amplifies extravasation at MTi and exacerbates the consequences of a loss of selectivity in endothelial transport, potentially exposing blood-borne molecules directly to neighbouring neurons usually located within 8-10 µm^20^.

**Figure 3.**
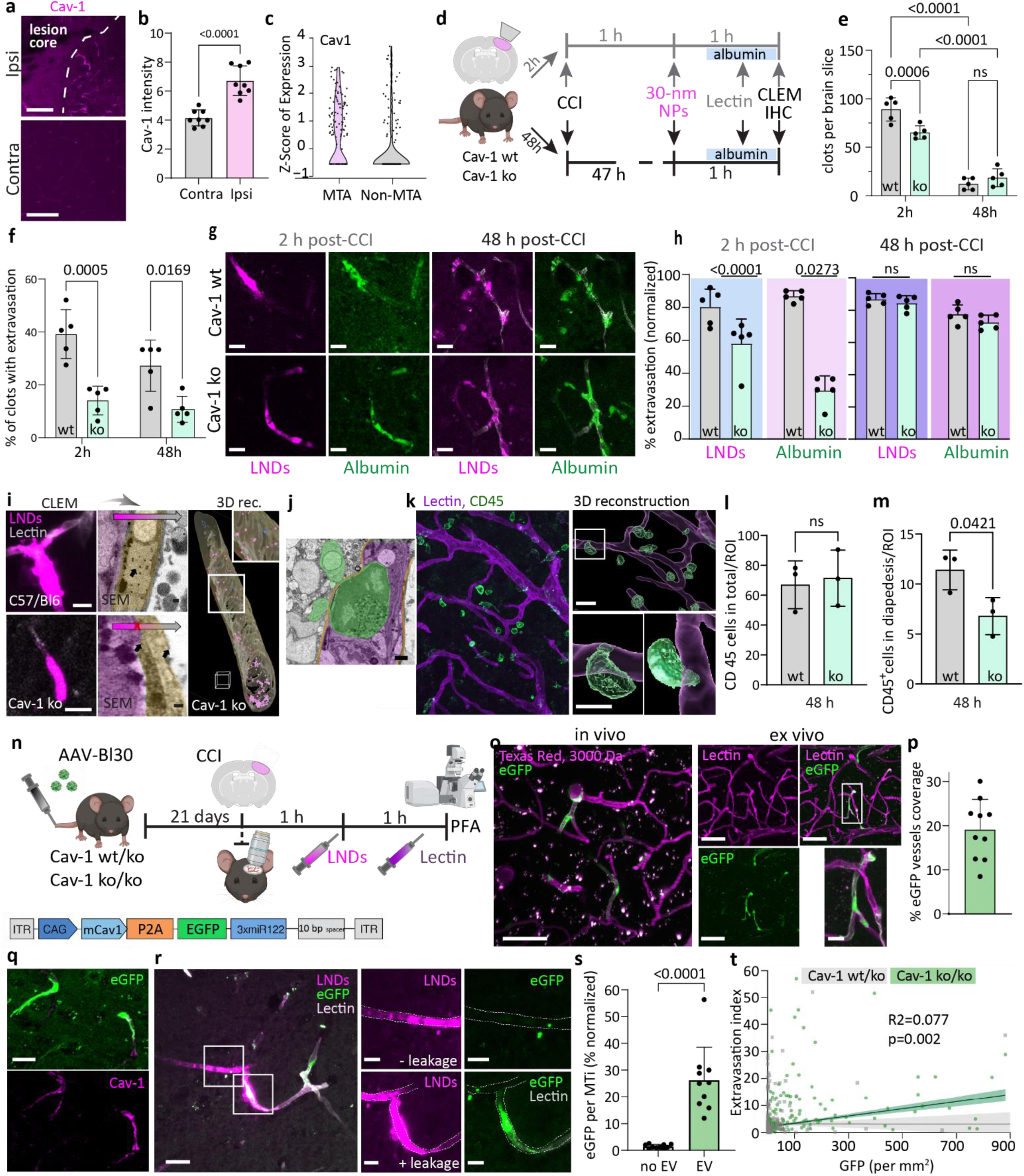
Cav-1 drives MTi-mediated BBB leakage and immune cell diapedesis after CCI. **a-b,** Representative confocal images and quantification of Cav-1 immunostaining in ipsilateral versus contralateral cortex 1 h post-CCI. Scale bar, 100 µm. Data are mean ± SD; n = 8 sections from 4 mice. **c,** Z-score analysis of BEC clusters from the dataset in Fig. 1e, showing enrichment of Cav-1 expression within the MTi-associated (MTA) endothelial cluster. **d,** Experimental design: Cav-1 wt and ko mice received 30-nm LNDs or gold NPs 1 h or 47 h after CCI, followed by albumin injection in 30 min and then lectin in 55 min, perfusion for IHC or CLEM. **e-f,** Quantification of LND-labelled MTi formation (e) and the proportion of MTi associated with BBB leakage (f) in Cav-1 wt and ko mice at 2 h and 48 h post-CCI. Data are mean ± SD; n = 5 mice per group; two-way ANOVA. **g-h,** Representative confocal images and quantification of MTi with LND or albumin extravasation (EV) in Cav-1 wt and ko mice at 2 h and 48 h post-CCI. Scale bar: 10 µm. Data are mean ± SD; n=5 mice, unpaired two-tailed t-test. **i,** Representative confocal and CLEM images (Original raw images are from Schifferer et al^9^.) showing MTi-mediated EV of NPs in C57BL/6N (upper panel) and no EV in Cav-1 ko mice (lower panel). 3D reconstruction (right panel) reveals NPs localization (magenta - intraluminal; blue – intracellular). Scale bars: confocal, 10 µm; SEM, 100 nm, 3D - 2 µm. SEM color overlay: MTi, magenta; endothelium, yellow; NPs, black. **j,** Representative SEM image showing a BEC (yellow), MTi (magenta) and immune cell (green) undergoing diapedesis. Scale bar, 2 µm. Original raw images are from Schifferer et al^9^. **k-m**, Representative confocal and 3D-reconstruction images and quantification of immune-cell diapedesis at 48 h post-CCI. Scale bars: confocal, 50 µm; 3D upper, 20 µm; 3D lower, 10 µm. Data are mean ± SD; n = 3 mice; 8 ROI per mouse. **n,** Experimental design of Cav-1 rescue study and virus construct. Cav-1 ko/ko and wt/ko mice were injected with AAV-BI30-Cav1-eGFP and analyzed 21 days later when endothelial Cav-1 expression was established. At this time point, mice underwent CCI (n = 5 per group), followed by LND injection 1 h post-injury and perfusion 1 h later for BBB imaging and Cav-1 restoration assessment. **o,** Representative two-photon *in vivo* and confocal *ex vivo* images showing cortical vasculature (*in vivo* - Texas Red–dextran, 3 kDa; *ex vivo* - lectin, magenta) and endothelial expression of AAV-BI30-Cav1-eGFP (green). Scale bars: 50 µm, zoomed – 10 µm. **p,** Quantification of virus transduction efficiency in BEC. Data are mean ± SD; n=10 mice (wt/ko+ko/ko). **q,** Representative confocal image showing co-localization of eGFP (AAV-BI30-Cav1-eGFP) and Cav-1 immunostaining in BECs. Scale bar: 20 µm. **r,** Representative confocal image from a Cav-1 ko mouse injected with AAV-BI30-Cav1-eGFP and subjected to CCI, showing two MTi: one with LND EV co-localized with eGFP signal and another without EV lacking eGFP expression. Scale bar, 20 µm; zoom, 10 µm. **s,** Quantification of eGFP fluorescence in MTi with versus without EV. Data are mean ± SD; n=10 mice. **t,** Regression analysis showing the relationship between LND extravasation index and eGFP signal density in Cav-1 rescue experiments. Each point represents an individual MTi (123 from Cav-1 wt/ko and 173 from Cav-1 ko/ko; 5 mice per group). A significant positive correlation was observed in Cav-1 ko/ko mice (green; R² = 0.077, p = 0.002), whereas no correlation was detected in Cav-1 wt/ko (grey; R² = 7.09 × 10⁻⁶, p = 0.9767).

We used original CLEM (Extended data Fig.4 a-g) to further investigate the ultrastructural profile of microthrombi-associated BECs. In C57BL/6N mice, CLEM from open data source (Schifferer et al, 2024^9^ from Kislinger et al., 2024^10^) confirmed the presence of numerous NPs within BEC at the site of MTi-BBB leakage (Fig. 3i, upper panel). In contrast, our CLEM in a Cav-1 ko mouse revealed a “bouton-like” LND-labelled MTi morphology without extravasation, a phenotype not previously observed in C57BL/6N animals (Fig. 3i, lower panel). After reconstruction of volume CLEM at single particle resolution we detected (Fig. 3i, right panel; Extended data Fig.4 g) that in Cav-1 deficient BECs, NPs accumulated intraluminally and are only present in negligible numbers within the cytoplasm, indicating that Cav-1 is indeed required for endothelial uptake of blood-borne cargo, and supporting the idea that MTi-BBB dysfunction is mediated by Cav-1–dependent transcytosis.

Finally, we assessed whether Cav-1 influences immune-cell trafficking. As noted above, we observed substantial recruitment of CD45⁺ cells at 48 h after TBI in both genotypes (Extended Data Fig. 3k–l and Extended Data Fig. 5a–b). Moreover, using our previously published open-access dataset (Schifferer et al, 2024^9^ from Kislinger et al.^10^), we demonstrate early leucocyte infiltration at sites of MTi (Fig. 3j). We therefore assessed immune-cell diapedesis from blood to brain after TBI and examined whether this process is modulated by Cav-1. At 2 h post-TBI, CD45⁺ cells were not detected within the parenchyma (data not shown). At 48 h however, substantial immune-cell infiltration could be observed, with CD45⁺ cells predominantly transmigrating through capillaries and additionally accumulating in the subarachnoid space (Extended Data Fig. 5a–b). Quantification of total parenchymal CD45⁺ cells revealed no differences across genotypes (Fig. 3l, Extended data Fig. 5c-d), indicating that the overall immune response to injury is comparable. In contrast, when specifically analyzing CD45⁺ cells undergoing diapedesis (Fig. 3k), we observe a marked reduction in Cav-1 ko animals (Fig. 3m, Extended data Fig. 5e-f), suggesting that Cav-1 supports capillary-associated immune-cell transmigration after TBI. Altogether, these findings establish Cav-1– dependent transcytosis as a major contributor to early MTi-associated BBB dysfunction after TBI, linking microvascular thrombi formation to altered molecular transport and cellular trafficking across the BBB.

### Endothelial Cav-1 re-expression rescues microthrombi-associated leakage

To directly test whether the Cav-1 dependence of MTi-associated BBB leakage originates from BECs, we generated an endothelial-tropic viral rescue system using the AAV-BI30 capsid^21^. This vector shows an exceptionally high specificity for BECs, and we packaged it with a Cav1–P2A–EGFP expression cassette, allowing simultaneous re-expression of Cav-1 and fluorescent identification of transduced endothelial cells. Intravenous delivery of AAV-BI30 into Cav-1 heterozygous (wt/ko) and Cav-1 homozygous (ko/ko) mice resulted in a mosaic population of Cav-1–positive (EGFP⁺) and Cav-1–negative (EGFP⁻) BECs within the same vascular network, providing an internal, vessel-level comparison of Cav-1 function under identical conditions (Fig. 3n). To determine the onset of endothelial expression of the viral construct *in vivo*, we performed longitudinal two-photon imaging, which revealed that EGFP fluorescence in BECs becomes robust and stable by day 21 after viral injection (Fig.3 o, in vivo). This defined the time point at which Cav-1 rescue could be reliably assessed. At 21 days post-AAV injection, we induced TBI and performed the same microthrombi-labelling paradigm used throughout the study: intravenous injection of 30-nm LNDs to visualize MTi and EV and lectin to label the vessel lumen. This enabled a direct comparison of MTi-associated extravasation between Cav-1–rescued (EGFP⁺) and Cav-1–deficient (EGFP⁻) capillaries within the same mouse. Using the contralateral hemisphere, confocal quantification showed that AAV-BI30 transduction covered ∼20% of the cortical microvasculature across all cohorts (Fig. 3o-p), and the eGFP signal colocalized with Cav-1 immunostaining, confirming successful endothelial Cav-1 re-expression in Cav-1 ko/ko mice (Fig. 3q). Interestingly, MTi without BBB leakage exhibited minimal green fluorescence (Fig. 3r), whereas MTi associated with BBB leakage displayed readily detectable and markedly stronger green signal (Fig. 3r), significantly exceeding that observed in non-leaking MTi (Fig. 3s). Finally, to determine whether BBB extravasation is indeed Cav-1-dependent, we performed a regression analysis relating Cav-1 expression to the magnitude of LNDs leakage. In Cav-1 heterozygous rescue mice (Cav-1 wt/ko), no meaningful correlation was observed, indicating that leakage events were essentially randomly associated with eGFP⁺ endothelial cells. In contrast, Cav-1 homozygous rescue mice (Cav-1 ko/ko) exhibited a significant positive correlation between Cav-1 expression (eGFP intensity) and LNDs extravasation (R² ≈ 0.077; p = 0.002), demonstrating that higher Cav-1 levels reliably associated with increased leakage (Fig. 3t). This confirms that Cav-1 is a key determinant of the extravasation mechanism at MTi sites.

### Microthrombus-driven brain damage is rescued by inhibiting leakage

To determine how Cav-1– dependent BBB impairment around microthrombi affects local NVU (neurovascular unit components), we compared microglial and neuronal responses at Cav-1 wt and ko mice at 2 h or 48 h after TBI (Fig. 4a). Importantly, quantification of the cortical penumbra (Fig. 4b) revealed that Cav-1 deficiency resulted in a significantly smaller penumbral area at 2 h post-CCI, whereas no genotype-dependent differences remained at 48 h, indicating an early but transient contribution of Cav-1 to tissue expansion after injury (Fig. 4c–d). Within the penumbral zone (Fig. 4b), correlative confocal microscopy showed that microglia rapidly accumulated around individual MTi already at 2 h after CCI and extended their processes towards LND-labelled clots (Fig. 4e). 3D reconstructions confirmed this microglia–microthrombus interface (Fig. 4f). Next, we analyzed microglial morphology in penumbral regions containing microthrombi (MTA) versus penumbral regions without microthrombi (non-MTA). In both genotypes, microglia in MTA regions exhibited significantly higher circularity and directionality compared with non-MTA microglia (Fig. 4g-h), suggesting specific additional local activation of microglia triggered by microthrombi-associated stimuli. However, in Cav-1 ko mice, which have substantially reduced BBB leakage, circularity of MTA microglia was significantly lower than in wt, while non-MTA microglia remained unchanged, suggesting that BBB dysfunction amplifies specific local activation of microglia which could be substantially attenuated when Cav-1-dependent transcytosis is blocked (Fig. 4g-h). This observation aligns with previous works showing that blood-borne components, like for instance fibrinogen, can exert potent pro-inflammatory effects on CNS-resident immune cells^22, 23^. Such findings support our interpretation that plasma proteins leaking through the BBB at microthrombus sites act as local inflammatory stimuli.

**Fig. 4.**
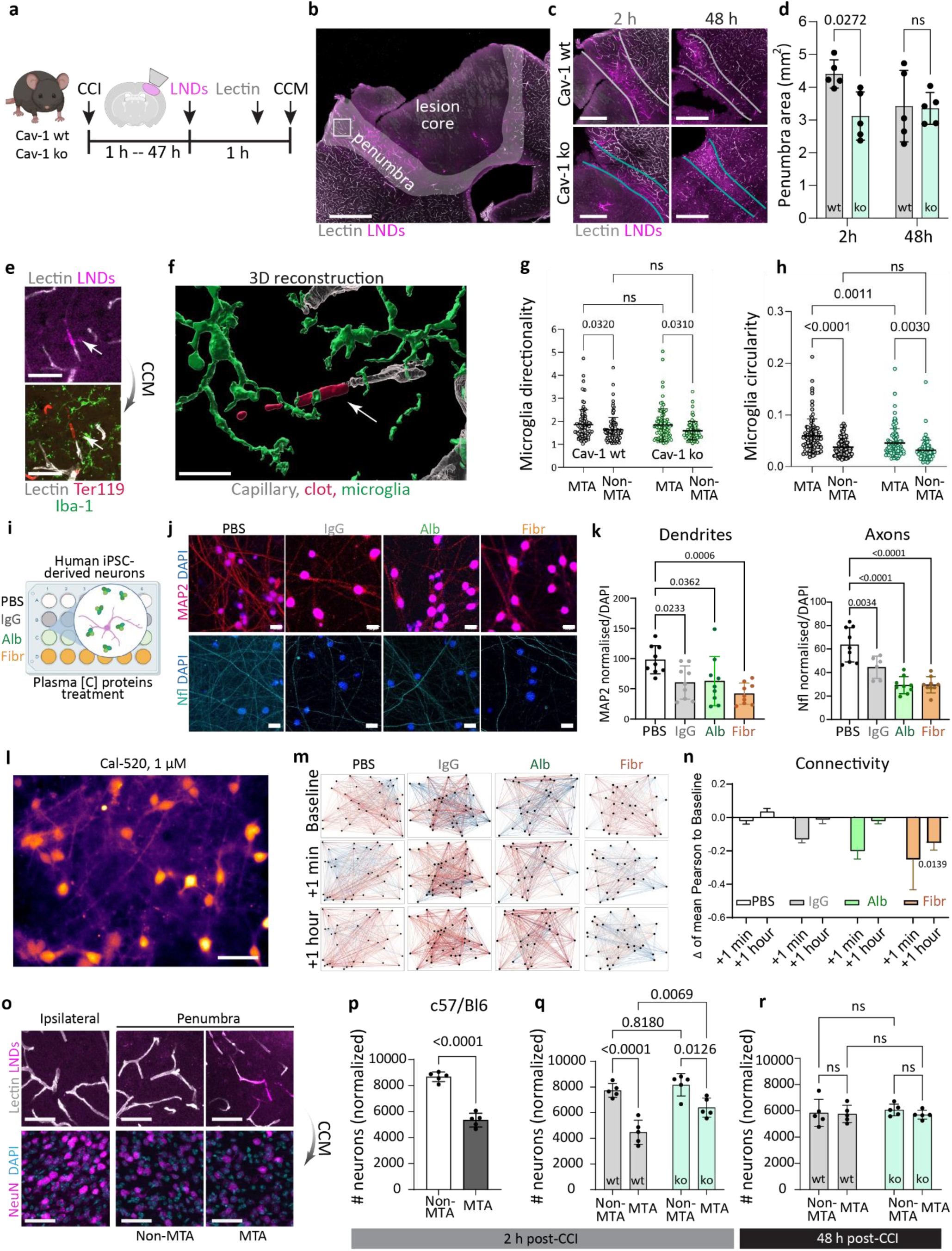
Microthrombus-associated Cav-1-dependent leakage promotes local neuroinflammation and neuronal damage. **a,** Experimental pipeline: Cav-1 WT and KO mice received 30 nm LNDs 1 h or 47 h after CCI, followed by lectin injection, perfusion, and tissue collection. **b,** Representative confocal image of lesion core and penumbra in Cav-1 WT. Scale bar: 500 µm. **c-d,** Representative images and analysis of penumbra area in Cav-1 WT and KO animals at different times post-CCI. Scale bar: 500 µm. Data are mean ± SD; n=5 mice, Two-way ANOVA. **e,** Correlative confocal microscopy (CCM) 1-hour post-CCI showing, in the penumbra (from panel b), microglia (iba-1, green) processes surrounding microthrombi (LNDs, Ter-119); Scale bar: 50 µm. **f,** 3D reconstruction of microglia-microthrombi interplay (see Movie S01). Scale bar: 20 µm. **g-h,** Analysis of microglia morphology in non-MTA (non–microthrombus-associated) *vs.* MTA (microthrombus-associated) regions: directionality (g); circularity (h). Data are mean ± SD; n=79-101 cells from 5 mice, Two-way ANOVA. **i,** Experimental pipeline: human iPSC-derived neurons (day 50 in vitro) were treated by PBS (control) or blood-borne proteins (IgG, albumin and fibrinogen) at 0.5x, 1x, 2x, human plasma-equivalent concentrations. After 24 hours, cells were fixed, MAP-2 immuno-stained and imaged. **j-k,** Representative images and analysis of iPSC-derived neurons (MAP-2) treated with plasma-equivalent concentrations of IgG, albumin, fibrinogen. Scale bar: 20 µm. Data are mean ± SD, n=9 ROI from 3 wells. One-way ANOVA. **l,** Representative Cal-520 fluorescence image of human iPSC-derived neuronal cultures used for calcium-imaging analysis. Scale bar: 100 µm. **m,** Representative functional connectivity maps generated from pairwise Pearson correlations of neuronal calcium activity at baseline, 1 min, and 1 h following exposure to PBS, IgG, albumin (Alb), or fibrinogen (Fibr). Nodes represent individual neurons and edges represent significant functional correlations between neuronal activity traces. **n,** Change in neuronal network connectivity quantified as the mean Pearson correlation coefficient relative to baseline (Δ). Data are mean ± SEM. n=3-4 independent dishes. Statistical analysis was performed using the Kruskal–Wallis test comparing each treatment with the PBS group. **o,** Representative CCM of capillaries, microthrombi, and NeuN-positive neurons in cortical ROIs (contralateral cortex; ipsilateral penumbra, non-MTA and MTA ROI) 2 h post-CCI. Scale bar: 50 µm. **p,** Analysis of neuronal death in non-MTA *vs.* MTA ROIs within the penumbra in C57Bl/6N mice, 2-hours post-CCI. Data are mean ± SD; n=5 mice, unpaired t-test. **q-r,** Analysis of neuronal death in Cav-1 WT and KO penumbral ROIs (non-MTA *vs.* MTA), 2 h and 48 h post-CCI. Data are mean ± SD; n=5 mice, Two-way ANOVA.

Since we demonstrated that extravasated blood components were able to reach local neurons, we next sought to determine whether blood-borne proteins exert direct neurotoxic effects. For this purpose, we used human iPSC-derived neurons and exposed them to plasma-equivalent concentrations of individual blood-borne proteins (Fig. 4i). After 24 h of exposure to albumin, IgG or fibrinogen we observed a significant dose-dependent reduction in both MAP-2 (dendrites) and Nfl (axons) signals, with albumin and fibrinogen exerting the strongest effects (Fig.4 j-k; Ext. data Fig. 6 a-b). Additionally, exposure to blood-borne proteins altered neuronal network connectivity hyper acutely, with fibrinogen producing the largest reduction in mean Pearson correlation, whereas albumin and IgG induced smaller effects (Fig. 4l–n; Extended Data Fig. 6c). Interestingly, this hierarchy repeats the kinetics of protein extravasation, suggesting that protein size may differentially influence transport and intracellular processing. Larger proteins such as fibrinogen may require larger transcytotic vesicles, slowing BBB passage, while simultaneously placing a greater burden on vesicle-mediated trafficking after uptake, thereby amplifying their impact on neuronal network function. Nevertheless, these *in vitro* findings indicate that even single blood-borne proteins may directly compromise both neuronal structure and function.

Finally, we examined whether MTi correlate with neuronal loss *in vivo*. In C57BL/6N mice 2 h post-CCI, NeuN quantification revealed significantly fewer neurons in MTA regions compared with adjacent non-MTA areas of the same penumbra (Fig. 4 o-p). The same pattern was observed in Cav-1 wt mice, indicating that MTi-associated leakage contributes to early neuronal loss (Fig. 4q). In contrast, Cav-1 ko mice exhibited a markedly reduced MTA-associated neuronal loss at 2 h consistent with reduced leakage of neurotoxic blood-borne components in Cav-1-deficient animals, while non-MTA associated neuron numbers stayed unchanged (Fig. 4q). However, these MTi-associated effects were not present at 48 h post-CCI (Fig. 4r), when, as was shown earlier, Cav-1-dependent transcytosis may not play a major role (Fig. 3h) and extravasated blood proteins predominantly accumulate in CD45+ immune cells (Extended Data Fig. 3 k-l).

In summary, these results indicate that MTi are local triggers of transient BBB impairment through upregulated, Cav-1-dependent, endothelial transcytosis which ultimately exposes neuronal and glial components to blood-borne components and thus promote local neuronal damage. Thus, targeting BBB leakage via Cav-1-dependent transcytosis may represent a promising therapeutic strategy to mitigate microthrombi-associated neurovascular impairment in brain injury.

### Clinical correlates of systemic coagulation dysregulation and vascular dysfunction after acute isolated TBI

To assess whether the coagulation and vascular alterations identified in our experimental models are reflected in human TBI, we integrated longitudinal blood parameters, neuroimaging and endothelial transcriptional profiles from patients with acute TBI. This multimodal approach enabled us to examine vascular alterations across complementary scales, from systemic haemostatic changes to cerebral vascular and endothelial abnormalities (Fig. 5; Extended Data Figs. 7, 8). We first characterized the haemostatic response to acute isolated TBI over time in patients without pre-injury anticoagulant (DOAC) or antiplatelet therapy. Acute TBI was associated with a rapid and dynamic haemostatic response, characterized by early reductions in circulating fibrinogen and platelet counts followed by distinct recovery trajectories (Fig. 5a). Fibrinogen displayed a particularly pronounced biphasic response, decreasing acutely before recovering and rising substantially above the healthy reference level over subsequent days. The broader longitudinal dataset additionally showed increased D-dimer together with changes in multiple coagulation parameters, consistent with acute coagulation activation and fibrin turnover (Extended Data Fig. 7d). A concomitant reduction in haemoglobin suggested that haemodilution and/or blood loss could contribute to the early changes, but did not account for the broader pattern of coagulation activation. We therefore focused on fibrinogen and analysed whether its early trajectory was modified by pre-injury antithrombotic treatment. Patients receiving DOAC or antiplatelet therapy showed higher circulating fibrinogen concentrations during the earliest post-injury period, with significant differences at 1–2 h and 6–7 h (Fig. 5b). Together with increased D-dimer and the early reduction in platelet counts, the relative preservation of circulating fibrinogen in antithrombotic-treated patients is consistent with reduced acute fibrinogen consumption and supports an early thrombotic/consumptive response following TBI. This observation was particularly relevant to our experimental findings, in which fibrin(ogen) constituted a major component of cerebral MTi and subsequently accumulated beyond the vascular compartment. Importantly, the effect of antithrombotic treatment on circulating fibrinogen was restricted to the acute post-injury phase, with the trajectories converging thereafter. Thus, the human haemostatic profile is consistent with a model in which acute fibrin formation and consumption accompany cerebral microvascular thrombosis, while BBB dysfunction enables extravasation of circulating fibrinogen into the injured brain. We next assessed whether early post-TBI coagulation abnormalities were associated with clinical outcome. Early coagulopathy was identified in 63 of 222 evaluable patients (28.4%; Fig. 5c). Patients with coagulopathy presented with lower admission GCS and higher Rotterdam CT scores and subsequently experienced longer ICU and hospital stays. Although median discharge GCS was 15 in both groups, the distribution was significantly shifted towards poorer neurological status in patients with coagulopathy (P=0.0084). Most notably, in-hospital mortality was approximately threefold higher in patients with coagulopathy than in those without coagulopathy (20.6% versus 6.9%, P=0.0068), whereas age and overall injury severity (ISS) did not differ significantly between groups. Together, these findings associate early post-TBI coagulopathy with greater neurological injury and an adverse clinical course. Additionally, neuroimaging revealed a spatial association between susceptibility abnormalities and FLAIR hyperintensity, supporting the coexistence of microvascular damage and surrounding parenchymal injury in acute TBI. (Fig. 5d; Extended Data Fig. 7a).

**Figure 5.**
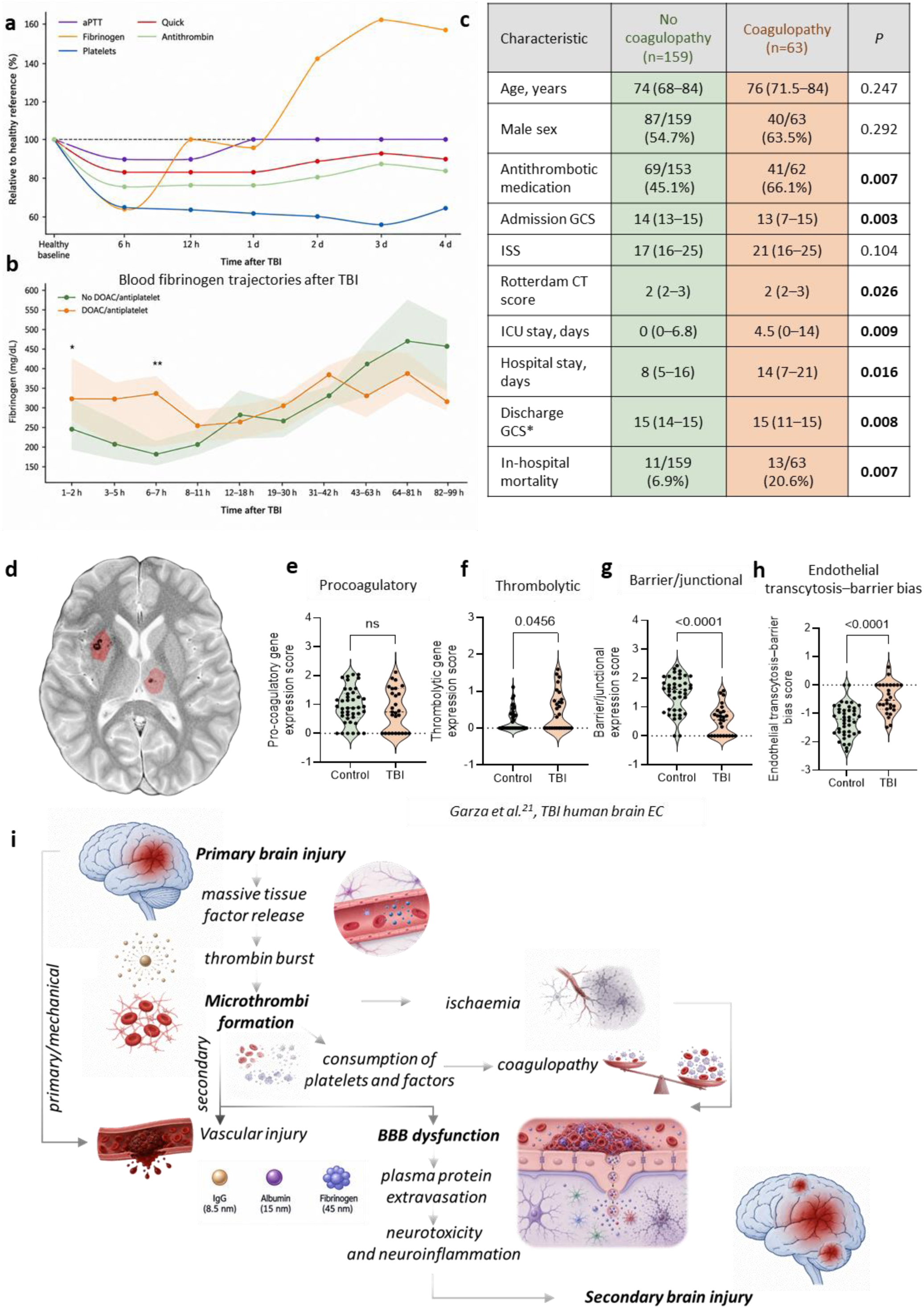
Human acute TBI exhibits coagulation dysregulation and vascular dysfunction. **a,** Clinical characteristics of patients with TBI stratified according to the presence or absence of coagulopathy (Early coagulopathy was defined a priori for this exploratory translational analysis as INR >1.2 and/or aPTT >35 s in any available sample obtained 1–11 h after admission.). Continuous variables are presented as median (IQR) and categorical variables as n/N (%). Bold P values indicate P < 0.05. GCS, Glasgow Coma Scale; ISS, Injury Severity Score; CT, computed tomography. Discharge GCS refers to GCS at hospital discharge. **b,** Temporal profiles of circulating haemostatic parameters after TBI in patients not receiving any anticoagulant (DOAC) or antiplatelet therapy. aPTT, Quick, platelet count, fibrinogen and antithrombin are expressed relative to the corresponding healthy reference value (100%) to illustrate their temporal trajectories from 6 h to 4 d after injury. The horizontal dashed line denotes the healthy reference level. **c,** Blood fibrinogen concentrations after TBI according to pre-injury DOAC/antiplatelet treatment. Lines indicate median fibrinogen concentrations and shaded regions indicate interquartile ranges across the indicated post-injury sampling windows. (* - P < 0.05; ** - P < 0.01, two-sided Mann–Whitney U test). **d**, Representative susceptibility-weighted imaging (SWI) from a patient with acute TBI, with FLAIR hyperintense signal overlaid in red (Extended Data Figure 7 a), illustrating the spatial relationship between traumatic tissue abnormalities and SWI-detected vascular injury. **e-h,** Analysis of the publicly available human brain RNA-seq dataset used to examine endothelial-cell (EC) transcriptional responses following TBI (Garza et al.^24^). Endothelial transcriptional signatures of pathway-level gene-expression scores of coagulation, transport, and barrier regulation after traumatic brain injury. Each dot represents one endothelial cell. Statistical significance was assessed using two-tailed Mann–Whitney tests. **i,** Proposed pathophysiological cascade linking primary TBI to secondary brain injury via cerebral microthrombi formation.

To determine whether these vascular abnormalities were accompanied by molecular alterations of the human BECs, we next re-analysed open data source of human TBI RNA-sequencing dataset (Garza et al., 2023)^24^, computing pathway-level scores from brain endothelial gene expression profiles. Given the limited number of human endothelial nuclei and the heterogeneity of injury timing and regions (Extended Data Fig.8a-d), we interpret these analyses as descriptive. While a composite pro-coagulatory endothelial gene score (VWF, EDN1, ANGPT2, ENG) was not significantly altered between control and TBI samples, TBI endothelial cells displayed a significant increase in a thrombolytic gene expression score (PLAT, ANXA2, ANXA11, SERPINE1). In parallel, a barrier/junctional gene expression score (CLDN5, PECAM1, TJP1, OCLN) was strongly reduced in TBI, indicating a loss of endothelial barrier identity. (Fig.5 e-h; Extended Data Fig.8e). Notably, a composite transcytosis–barrier score revealed a shift towards caveolae-associated transcytotic programs in TBI endothelial cells compared with controls (Fig. 5h), providing an independent human correlate of the Cav-1-associated increase in BBB transcytosis observed experimentally.

Finally, we integrated these findings into a model linking coagulation dysregulation and vascular dysfunction in acute TBI (Fig. 5i). Primary brain injury and massive tissue factor release promote thrombin generation and fibrin-rich cerebral microthrombus formation, accompanied by consumption of circulating haemostatic factors. Microvascular thrombosis and associated ischaemic injury may further compromise BBB function, facilitating transendothelial extravasation of circulating proteins accumulated inside MTi, including fibrinogen, into the brain parenchyma. Together with the direct neurotoxic effects of extravasated plasma proteins identified experimentally, these processes provide a potential link between acute coagulation activation, BBB dysfunction and secondary brain injury.

## Discussion

TBI represents a major and growing global health burden and is one of the most prevalent neurological disorders worldwide, with increasing incidence^25^. Disturbances of haemostasis are a prominent feature of acute TBI and are associated with injury severity and poor clinical outcome^26^. Within the injured brain, activation of coagulation can result in widespread cerebral microthrombosis, and neuropathological studies some decades ago have demonstrated a strong association between intravascular microthrombi and selective neuronal necrosis^27^. Despite this association, the mechanisms linking post-traumatic microthrombosis to progressive secondary neuronal injury remain incompletely understood.

Here, we identified BBB dysfunction as a missing link and critical component of this process. Namely, early BBB permeability is a focal and spatially organized phenomenon. Highly fluorescent nanoscale probes revealed leakage sites confined to individual microvessels within the penumbra. These focal permeability sites consistently colocalized with microthrombi. Ultrastructural analyses further showed that tracer extravasation occurred via morphologically intact endothelial cells adjacent to MTi. Importantly, not all MTi were associated with detectable leakage, suggesting that BBB dysfunction is not an inevitable consequence of microvascular occlusion but instead reflects a context-dependent endothelial response. Together, these findings redefine early BBB dysfunction after TBI as a microthrombus-linked, focal event rather than a diffuse consequence of global vascular injury.

Closer examination of these microthrombus-associated leakage sites revealed that BBB dysfunction is not a passive loss of barrier integrity, but a regulated endothelial process with clear size selectivity. Blood-borne components and nanoparticles first stop their motion, accumulate within microthrombi and then cross the adjacent endothelium before dispersing into the surrounding parenchyma, indicating a directed transport process. These observations suggest that microthrombi act both as local reservoirs of circulating factors and as triggers of focal transendothelial transport. The kinetics and spatial spread of extravasation varied markedly between molecules of different sizes, with smaller proteins penetrating more rapidly and over greater distances than larger plasma components. The upper size is probably limited by size of BEC caveolae, which is around 50 nm. Moreover, the presence or absence of leakage varied between individual microthrombi and correlated with their cellular composition, particularly the recruitment of immune cells and matrix-remodeling activity. These observations argue against a uniform breakdown of the BBB and instead support a model in which endothelial transport pathways are locally engaged in response to microthrombus-associated cues. This spatially restricted and size-dependent permeability provides a plausible mechanism by which circulating factors gain focal access to neural tissue during the early phase of secondary injury.

We found that caveolae-mediated endothelial transport pathways are involved in the MTi-associated increase in permeability. It is widely known that suppression of caveolae-mediated transcytosis is a defining feature of the matured intact BBB and its local reactivation represents a regulated pathological switch^28^. A recent study in SARS-CoV-2 demonstrates that increased endothelial Cav-1 is sufficient to promote transcellular permeability, neuroinflammation, and cognitive impairment, while genetic Cav-1 deficiency confers vascular and functional protection^18^. Our findings show that Cav-1–dependent transcytosis is locally and transiently active at sites of MT formation, producing focal BBB dysfunction without widespread barrier collapse. Genetic deletion of Cav-1 markedly reduced leakage of both endogenous plasma proteins and LNDs at MT-containing vessels, indicating that transcellular transport without junctional disruption dominates this early phase of BBB dysfunction. Notably, Cav-1 deficiency did not simply attenuate transport of canonical caveolar cargo such as albumin, but also limited the extravasation of larger nanoparticles, suggesting a loss of molecular selectivity when caveolar pathways are pathologically engaged. This Cav-1–dependent permeability is temporally restricted but BBB leakage reappears 48 h post-CCI when junctional alterations become more prominent, implicating a dynamic shift in BBB transport mechanisms over time. Similar biphasic BBB permeability changes have been shown acutely after stroke and TBI^19, 29^. Together, these observations place caveolae-mediated transcytosis at the center of early, MT-associated BBB dysfunction after TBI. By reconstituting endothelial Cav-1 expression in Cav-1–deficient mice through viral transfection, we could selectively restore microthrombus-associated leakage in our target vessels, showing that Cav-1 acts locally within the endothelium to control permeability. Importantly, Cav-1 deficiency reduces extravasation and edema, leading to preservation of the capillary lumen. These combined effects may also indirectly contribute to relative reduction of MTi formation.

The focal nature of microthrombus-associated BBB dysfunction also acts on the surrounding parenchyma. Microglial processes rapidly expand towards MT-containing vessels, consistent with previous reports of microglial response to vascular injury^30, 31^. Suppression of Cav-1–dependent transcytosis reduced, but did not eliminate microglial activation, suggesting that blood-borne components amplify this response while additional microthrombus-or endothelium-derived cues also may contribute, including ATP/ADP-driven, P2Y12-dependent microglial chemotaxis triggered by vascular and tissue injury^32^. Further, Cav-1-dependent transcytosis may enable entry of specific size-fitting blood-borne components such as fibrinogen. With its large size and high local concentration within microthrombi, it has been shown to trigger rapid perivascular microglial responses^33^. Recent works in a cerebral amyloid angiopathy model associate fibrinogen extravasation with increased endothelial Cav-1 and microglial activation, showing that fibrinogen depletion reduces Cav-1 levels, dampens microglial activation, and improves memory function^34^. Consistent with these findings, we demonstrate that extravasated plasma proteins can reach nearby neurons. Exposure of human neurons to physiologically relevant concentrations of blood-derived proteins induces neuronal structural damage and alters neuronal activity and network connectivity. These functional effects also show a size-dependent pattern, with fibrinogen producing the most pronounced impairment, suggesting that protein size may influence neuronal endocytosis, although fibrinogen-specific signalling is also likely to contribute. *In vivo*, neuronal loss was selectively observed in MT-associated regions and was markedly reduced in the absence of Cav-1-dependent leakage. These findings indicate that focally increased BBB permeability around microthrombi creates a specific microenvironment, promoting neuroinflammation and neuronal injury and thus links early vascular dysfunction to localized neurodegenerative changes after TBI. This mechanism is further supported by our clinical observations, in which the acute decrease in circulating fibrinogen was attenuated by pre-injury antithrombotic treatment and restricted to the early post-injury phase, consistent with transient fibrinogen consumption during acute MTi formation and in line with other studies^35^. The timing of BBB dysfunction appears critical, as early microthrombus-associated leakage occurs before substantial immune cell recruitment, allowing large blood-borne molecules to directly access neural tissue. At later stages, permeability may rather be dominated by paracellular transport that are unlikely to permit the passage of large molecules, and extravasated components are massively captured by infiltrating immune cells, indicating a fundamentally different and potentially less harmful mechanism.

Clinical longitudinal blood profiling revealed a dynamic haemostatic response to acute TBI, characterized by early coagulation activation and consumption followed by distinct recovery trajectories, while early coagulopathy was associated with greater neurological injury and increased mortality. The coexistence of dynamic coagulation dysregulation, focal vascular abnormalities on neuroimaging, and an endothelial transcriptional shift away from junctional barrier identity towards caveolae-associated transport suggests that the vascular response to acute TBI extends across systemic and cerebral compartments. Importantly, the association of early coagulopathy with poorer neurological status and mortality indicates that these haemostatic alterations are associated with clinically meaningful consequences. Moreover, other SWI– pathology correlation studies indicate that detectable traumatic microbleeds are usually accompanied by additional microvascular injuries downstream of damaged vessels that fall below the resolution of SWI, but can be observed histologically and are often surrounded by activated microglia.^36^ This is particularly relevant to our observation that BBB leakage is confined to individual microvessels: the absence of a detectable imaging abnormality may therefore not imply an intact microvascular environment. Clinically visible lesions may represent only a fraction of a substantially larger burden of focal vascular dysfunction. Together, these findings suggest that focal microvascular events can initiate highly localized endothelial, inflammatory and neuronal responses that extend beyond the vascular lesion itself. Although individual lesions may remain clinically silent, their cumulative burden could contribute to progressive secondary injury and potentially to the long-term neurological consequences of TBI.

From a therapeutic perspective, our findings highlight a narrow but potentially actionable window during which MT-associated BBB dysfunction is mediated by regulated endothelial transport before the emergence of more extensive structural barrier alterations. The Cav-1-dependent nature of this early permeability, together with the ability to restore leakage through endothelial Cav-1 re-expression, indicates that this process is reversible and locally controlled at the level of the endothelium. These observations suggest that selective modulation of endothelial transport pathways could mitigate early secondary injury while preserving vascular homeostasis and avoiding the broader consequences of systemic suppression of coagulation or inflammation.

At the same time, several limitations of our study should be acknowledged. Our mechanistic analyses were performed primarily in experimental models, and although supported by convergent clinical imaging, blood-based, and transcriptional data, direct visualization of MTi and BBB transport mechanisms in the human brain remains beyond current clinical technologies. In addition, our study focuses on the acute stage of TBI and does not address long-term cognitive outcomes. Future work will be required to define how early, focal vascular events interact with chronic vascular stress and to determine whether modulating endothelial transport can alter long-term neurological trajectories.

## Conclusion

We identify cerebral microthrombi as focal points of early, Cav-1-dependent, size-selective BBB dysfunction that locally exposes neural tissue to blood-borne components, triggering neuroinflammation and neuronal damage after traumatic brain injury.

## Material and Methods

### Experiment Design

All animal experiments were approved by the Ethical Review Board of the Government of Upper Bavaria and conducted in accordance with the ARRIVE guidelines. Animal husbandry, health monitoring, and hygiene management were performed in accordance with the recommendations of the Federation of European Laboratory Animal Science Associations (FELASA). Investigators were blinded during image acquisition and data analysis. For experiments characterizing microthrombosis and blood–brain barrier dysfunction, animals were randomly allocated to the respective injection groups (saline versus LnDs; albumin versus fibrinogen versus IgG). For the mechanistic experiments, Cav1 wild-type, knockout, and heterozygous animals were randomized by an independent researcher who was not involved in the experimental procedures or subsequent analyses. Investigators performing the experiments and analyses remained blinded to genotype throughout the study. A total of 44 animals were used for this project. For the characterization of microthrombosis, 19 male C57BL/6N mice weighing 28–33 g underwent controlled cortical impact (CCI) as described below. Sixty minutes after CCI, mice received 30 nm LNDs via an intra-arterial catheter. To assess BBB disruption, animals received intra-arterial injections of fluorescently labeled albumin, fibrinogen, or IgG. For caveolin pathway experiments, Cav1 knockout (n = 5), Cav1 wild-type (n = 5), and heterozygous Cav1 mice (n = 5) of mixed sex underwent CCI using the same protocol. For viral reconstitution experiments, Cav1 knockout (n = 5) and heterozygous mice (n = 5), also of mixed sex, were treated identically. To visualize perfused cerebral vessels, DyLight 649–labeled Lycopersicon esculentum lectin was administered 5 min before sacrifice. Animals were deeply anesthetized and transcardially perfused with 4% paraformaldehyde (PFA) 120 min after traumatic brain injury. Brains were removed and 50 μm-thick coronal sections acquired through the lesion region using a vibratome, as previously described. Three representative sections per animal were mounted in Fluoromount, coverslipped, and imaged by confocal microscopy.

### Controlled Cortical Impact

Experimental traumatic brain injury was induced using the controlled cortical impact model as previously described^14^. Briefly, animals received preoperative analgesia with buprenorphine (0.05 mg/kg) 30 minutes before surgery. Anesthesia was induced with isoflurane (5%) and maintained with 1.5–2.5% isoflurane in an oxygen/air mixture under continuous monitoring of body temperature and heart rate. Following right parietal craniotomy, impact was delivered directly onto the intact dura using a pressure-controlled custom-built CCI device (L. Kopacz, University of Mainz, Germany) at a velocity of 6 m s⁻¹, penetration depth of 1 mm, and contact time of 150 ms. After impact, the skull plate was repositioned and the wound was closed. Animals were maintained in an incubator at 34 °C and 60% humidity to prevent hypothermia until recovery/sacrificing.

### Lipid nanodroplets (LNDs) formulation

60 mg of Kolliphor® ELP, and 40 mg of LabrafacTM (with 1 wt.% R18-TPB) are mixed with a stirring bar in an Eppendorf, at 60 °C for 5 min. While stirring, filtered MQ water at 60 °C is added in one shot. The mixture is vortexed for 30 sec, and homogenized in a thermoshaker for 10 min (60 °C, 1400 rpm). LND’s diameters are determined by dynamic light scattering (DLS) on a Zetasizer Nano (Malvern).

### Catheterization and Injection of Fluorescent Markers

Fluorescent tracers and nanoparticles were administered through femoral arterial catheterization using an established protocol. Animals were anesthetized with medetomidine, midazolam, and fentanyl and placed in the supine position on a heating pad. The hind limb was fixed, fur was removed, and the surgical field was disinfected. A longitudinal skin incision was made to expose the femoral neurovascular bundle. The femoral artery was carefully separated from the vein and nerve, and a latex strip was placed underneath the artery to facilitate visualization. A small lateral arteriotomy was made using microsurgical scissors, the vessel was gently dilated, and the catheter was inserted and advanced approximately 1–2 cm. Correct placement was confirmed by blood inflow into the catheter, after which the catheter was secured with ligature. The wound was sealed with tissue adhesive and the animal was placed into a heated recovery chamber. LNDs were diluted 1:25 and injected at 4 μL g⁻¹ body weight. DyLight 649–labeled Lycopersicon esculentum lectin (Vector Laboratories, Burlingame, CA, USA; 1 mg mL⁻¹) was injected at a volume of 0.1 mL per mouse for vascular labeling. Fluorescent plasma protein tracers were administered at concentrations approximating physiological plasma levels: IgG at 10 mg mL⁻¹ (Alexa Fluor Plus 488 conjugate, Thermo Fisher Scientific, A32814), fibrinogen at 2 mg mL⁻¹ (Alexa Fluor 488 conjugate, Thermo Fisher Scientific, F13191), and albumin at 5 g dL⁻¹ (Alexa Fluor 488 conjugate, Thermo Fisher Scientific, A13100), each at 0.1 mL per mouse. For correlative light and electron microscopy, 30 nm PEG2000-coated, methyl-terminated gold nanoparticles (density 1.00 g cm⁻³; Sigma-Aldrich, St. Louis, MO, USA) were injected through the femoral catheter at the same dose used for LNDs.

### Confocal Image Acquisition

Confocal imaging was performed using a ZEISS LSM 980 microscope (Carl Zeiss Microscopy GmbH, Jena, Germany). Images were acquired with a 20× objective (EC Plan-Neofluar 20×/0.50 Pol M27) using an image matrix of 512 × 512 pixels, corresponding to a pixel size of 0.4 × 0.4 μm and 8-bit depth. Sections were first imaged prior to staining to capture intrinsic nanoparticle fluorescence, then recovered as free-floating sections, immunostained, and reimaged under identical conditions. Three brain sections per animal were analyzed for each staining condition. For high-resolution imaging of the lesion area, z-stacks spanning 50 μm were acquired at 2 μm intervals, covering the lesion core and surrounding penumbra. Whole-lesion overviews were obtained as tile scans with a slice interval of 5 μm over a total depth of 50 μm.

### Immunohistochemistry

Animals were deeply anesthetized and transcardially perfused with 4% PFA. Coronal free-floating sections (50 μm) were prepared using a vibratome. Sections were permeabilized with 0.1% Tween-20 in PBS for 1 h at room temperature and then incubated for 48 h at 4 °C in antibody buffer containing 1% bovine serum albumin, 0.1% cold-water fish skin gelatin, and 0.5% Triton X-100 in 0.01 M PBS, pH 7.2–7.4, together with the primary antibodies. The following primary antibodies were used: TER119 (rat, Abcam, ab91113, 1:200), fibrin (mouse, GeneTex, GTX19079, 1:100), Ly6C/G (rabbit, MyBioSource, MBS488606, 1:100), caveolin-1 (rabbit, Cell Signaling Technology, 3238, 1:100), NeuN (rabbit, Abcam, ab177487, 1:200), Iba-1 (rabbit, WAKO, 019-19741, 1:200), ZO-1 (rabbit, Invitrogen, 61-7300, 1:100), and claudin-5 (mouse, Thermo Fisher Scientific, 35-2500, 1:50). After washing in PBS, sections were incubated with species-specific secondary antibodies diluted 1:200 in 0.05% Tween-20 in 0.01 M PBS. Secondary antibodies included Alexa Fluor 594– conjugated goat anti-rabbit (Thermo Fisher Scientific, A-11012), Alexa Fluor 488–conjugated goat anti-rat (Thermo Fisher Scientific, A-11006), Alexa Fluor 594–conjugated goat anti-mouse (Thermo Fisher Scientific, A-11005), and Alexa Fluor 488–conjugated goat anti-mouse (Thermo Fisher Scientific, A-11001). Nuclei were counterstained with DAPI (Invitrogen, D1306; 1:10,000 in PBS).

### Correlative Fluorescence Microscopy

For correlative fluorescence microscopy, fixed brains were sectioned coronally at 50 μm thickness using a vibratome. Three representative sections per brain were mounted in Fluoromount diluted 1:10 in PBS to permit subsequent section recovery, covered with glass coverslips, and imaged by confocal microscopy. Sections were initially imaged without immunostaining to record intrinsic LND and lectin signals. Slides were then immersed in PBS for 1 h to allow removal of the coverslip, and sections were gently detached and washed overnight in PBS before undergoing immunohistochemistry. Lectin labeling of perfused vessels served as an anatomical reference to relocate identical regions of interest in pre-and post-staining images of the same section.

### Analysis of Clot Formation

To assess the effect of LNDs on microvascular thrombus formation, sections from animals injected with LnDs or PBS (n = 5 per group) were analyzed using correlative fluorescence microscopy. For quantification of clot number, pre-stained sections were analyzed in Fiji/ImageJ. Binary masks were generated from areas showing overlap between lectin and LnD signals, and the number of particles was quantified using the “Analyze Particles” function.

For clot composition analysis, sections were stained for TER119 and fibrin. Raw confocal images were imported into Fiji and channels were separated. The penumbra was defined as the lectin-positive vascular territory containing erythrocyte and/or fibrin accumulation within vessels. Hemorrhagic areas with erythrocytes located outside the vascular lumen were excluded. The penumbra was cropped in all channels, after which the lectin channel was thresholded and binarized to generate a vascular mask. Skeleton analysis was performed using the “Analyze Skeleton” plugin in Fiji. To identify erythrocyte stalls, fibrin-containing occlusions, and thrombi, channel overlap images were generated using the “Image Calculator” function. Logical overlap of TER119 with lectin identified erythrocyte stalls, overlap of fibrin with lectin identified fibrin-positive occlusions, and overlap of both resulted in thrombi. These images were thresholded, skeletonized, and quantified for number of occlusions, branch number, vessel diameter, and total vessel length. Values were normalized to the total lectin-positive vessel length within each image.

### Analysis of LND and Albumin Extravasation

The lesion penumbra was used as the region of interest for extravasation analysis. Mean fluorescence intensities of lectin, LNDs, and albumin were quantified in Fiji. Colocalization between lectin and LND or albumin signals was assessed using the JACoP plugin with identical threshold settings across all samples. Colocalized signal was interpreted as intravascular tracer, whereas the non-colocalized fraction represented extravasated material.

The border between perfused and non-perfused tissue was defined based on DyLight 649 lectin labeling. This border was measured in Fiji and designated as the lesion perimeter. The area of tracer extravasation extending into adjacent tissue outside the perfused border was quantified and normalized to lesion perimeter length to calculate the average depth of extravasation. Extravasation intensity was determined as the mean gray value in the extravasated area. Measurements were performed on three coronal sections per animal.

### Characterization of Clot Composition

Fluorescence intensity within vascular occlusions was quantified in Fiji as mean gray value within manually defined ROIs. Three coronal sections per animal were analyzed, yielding a total of 146 occlusions in the 30 nm group and 341 occlusions in the 80 nm group. Raw ZEISS image files were imported into ImageJ, converted to maximum-intensity projections, and analyzed in defined ROIs. Microglial coverage of occlusions was assessed by manual counting of Iba-1-positive cells associated with clot-containing vessels.

### Correlative light-electron microscopy

CLEM was performed following a recently published workflow^10, 14^. Briefly, mice were perfused with fixative containing 4% paraformaldehyde and 2.5% glutaraldehyde in 0.1 M sodium cacodylate buffer (pH 7.4), and dissected brains were immersion-fixed for 24 h. Coronal vibratome sections were prepared and post-fixed for an additional 24 h before storage in PBS. Regions of interest were identified by confocal imaging and trimmed accordingly. Samples were processed using a standard rOTO en bloc staining protocol^37^, including osmium tetroxide, thiocarbohydrazide, and uranyl acetate contrasting, followed by dehydration and resin embedding (LX112). Serial ultrathin sections (∼100 nm) were collected on carbon nanotube tape using an automated tape-collecting ultramicrotome and mounted on silicon wafers. Electron microscopy imaging was performed using a scanning electron microscope with backscatter detection. Large-scale volumes were acquired to enable correlation with confocal datasets based on vascular landmarks and anatomical features. Regions containing microthrombi were subsequently imaged at higher resolution. Image stacks were aligned and processed using Fiji/TrakEM2. Manual segmentation was performed in webknossos (scalable minds) and, subsequent rendering was done in Blender^38^.

### Endothelial Cell Isolation

Mice were anesthetized and transcardially perfused with cold saline. Brains were collected, washed in cold Dulbecco’s PBS (D-PBS; Gibco), and cut into eight sagittal slices on ice. Tissue was enzymatically and mechanically dissociated in gentleMACS C Tubes using Adult Brain Dissociation Kit enzymes and a gentleMACS Octo Dissociator. Dissociated tissue was filtered through 70 μm MACS SmartStrainers (Miltenyi Biotec), washed, and centrifuged at 300 × g for 10 min at 4 °C. Myelin was removed using Myelin Removal Beads II and MACS LS Columns (Miltenyi Biotec). CD45⁺ cells were then depleted using CD45 MicroBeads and LD Columns, followed by enrichment of CD31⁺ endothelial cells using CD31 MicroBeads and LS Columns (Miltenyi Biotec). Endothelial enrichment was confirmed by flow cytometry. The isolated fraction was stained with a fluorophore-conjugated anti-CD31 antibody, washed by centrifugation, and resuspended in D-PBS/BSA buffer before flow cytometric analysis.

### CD45 Quantification

For assessment of leukocyte depletion during endothelial enrichment, cell suspensions were subjected to magnetic depletion of CD45⁺ cells using CD45 MicroBeads and LD Columns as described above. The resulting fractions were evaluated by flow cytometry in combination with CD31 staining to confirm depletion of hematopoietic cells and enrichment of endothelial cells. CD45 depletion efficiency was assessed on the basis of the relative abundance of CD45-positive and CD31-positive populations in the isolated cell fraction.

### Adeno-Associated Virus Production, Purification and Titration

AAV vectors were generated with assistance from VectorBuilder. The plasmid pAAV-CAG-BI30-Cav1-eGFP-WPRE was packaged using the BI30 capsid AAV9 by polyethyleneimine-mediated triple transfection of HEK293-T17 cells with the transfer plasmid, the BI30 rep/cap packaging plasmid, and pHelper plasmid. Viral particles were harvested 72–96 h after transfection. Virus was precipitated from cell lysates and culture medium using polyethylene glycol 8000, collected by centrifugation, and further processed by chloroform extraction followed by aqueous two-phase extraction with ammonium sulfate and polyethylene glycol 8000. Additional purification was performed by discontinuous iodixanol gradient ultracentrifugation, and virus was concentrated using centrifugal filter columns.

Viral genome titers were determined by quantitative PCR targeting eGFP. Extracapsid DNA was removed by DNase I digestion, after which encapsidated viral DNA was released by alkaline lysis. Quantitative PCR was performed using TaqMan Fast Advanced Master Mix (Applied Biosystems) and the following GFP-specific oligonucleotides: forward primer 5′-GAACCGCATCGAGCTGAA-3′, reverse primer 5′-TGCTTGTCGGCCATGATATAG-3′, and probe 5′-/56-FAM/ATCGACTTC/ZEN/AAGGAGGACGGCAAC/3IABkFQ/-3′. Final titers exceeded 10¹³ genome copies per mL.

### AAV Application, Imaging and Analysis

BI30-AAV particles were administered by tail vein injection at a dose of 1.5 × 10¹² vector genomes in 100 μL per mouse. Cav1 knockout and heterozygous mice were injected. To determine the optimal expression interval, one animal was imaged weekly after injection using two-photon microscopy. Peak expression was observed 21 d after injection, and this interval was therefore used for subsequent traumatic brain injury experiments.

Two hours after CCI, animals underwent lectin and LND administration followed by perfusion and brain collection as described above. Confocal microscopy was used to acquire eGFP, LND, and lectin signals without additional immunostaining. Fluorescence channels were separated in ImageJ, and ROIs corresponding to microvascular clots were manually selected on the basis of LND signal. Ten ROIs per section were analyzed across three sections per animal, including clots with and without evidence of extravasation. An extravasation index was calculated as the ratio of LND signal located outside lectin-positive vessels relative to total LND signal area. eGFP signal was quantified separately within the same ROIs, and the relationship between Cav1 expression and the extravasation index was assessed using Pearson correlation in GraphPad Prism.

### Human iPSC Culture and Cortical Differentiation

The human induced pluripotent stem cell line hiN.Fm.m.LON71-019 (passages 28–31) was obtained from the Nantes iPSC Core Facility and routinely tested for mycoplasma contamination. Cells were maintained under feeder-free conditions on Cultrex-coated plates in mTeSR medium (STEMCELL Technologies) at 37 °C in a humidified atmosphere containing 5% CO₂. Medium was replaced daily. Cells were passaged every 4–5 d using Versene (0.48 mM EDTA; Gibco) as was described earlier^39^.

Cortical differentiation was performed as previously described. For neural induction, hiPSC colonies were transferred to N2B27 medium consisting of 50% DMEM/F-12 GlutaMAX, 50% Neurobasal medium, 2% B27 supplement without vitamin A, 1% N2 supplement, and 50 μM β-mercaptoethanol. Cells were incubated for 6 h in low-adhesion plates in N2B27 supplemented with SB431542 (20 μM), LDN-193189 (0.1 μM), and Y-27632 (10 μM). Following neural aggregate formation, cells were transferred to plates coated with poly-L-ornithine (15 μg mL⁻¹) and laminin (10 μg mL⁻¹).

From day in vitro (DIV) 0 to DIV20, cultures were maintained with daily medium changes. Y-27632 was withdrawn at DIV1. Canonical WNT signaling was inhibited with XAV-939 (1 μM) from DIV1 to DIV9. Between DIV5 and DIV9, SB431542 was withdrawn and FGF2 (10 ng mL⁻¹) together with cyclopamine (1 μM) was added to promote dorsal telencephalic fate. From DIV10 to DIV17, LDN-193189 and XAV-939 were removed and WNT signaling was mildly activated using CHIR99021 (0.4 μM) to support neuronal progenitor maturation.

At DIV17, cortical progenitors were dissociated with Accutase for 5–7 min at 37 °C, centrifuged at 200 × g for 5 min, resuspended at 5 × 10⁶ cells mL⁻¹ in CryoStor cryopreservation medium, and frozen using a controlled-rate freezing container before long-term storage in liquid nitrogen at −150 °C. For terminal differentiation, cryopreserved progenitors were thawed, washed, and plated on poly-L-ornithine- and laminin-coated plates at approximately 100,000 cells cm⁻². Cells were initially maintained in N2B27 medium supplemented with BDNF (20 ng mL⁻¹), dbcAMP (100 μM), DAPT (10 μM), and Y-27632 (10 μM). Twenty-four hours later, Y-27632 was replaced by PD0332991 (2.5 μM). Thereafter, half of the medium was replaced weekly with BrainPhys Neuronal Medium supplemented with SM1 and N2A. DAPT was progressively reduced over subsequent weeks, and PD0332991 was similarly tapered before withdrawal.

### Protein Treatments

For immunocytochemistry experiments, cortical neurons were plated at 100,000 cells cm⁻² in poly-L-ornithine- and laminin-coated 96-well plates. After 6 weeks of terminal differentiation, neurons were exposed for 24 h to human fibrinogen (Sigma-Aldrich, F3879; 1, 4, or 8 g L⁻¹), human serum albumin (Sigma-Aldrich, A1653; 17, 50, or 100 g L⁻¹), or human IgG (Sigma-Aldrich, 56834; 3, 15, or 30 g L⁻¹). Each condition was tested in three independent wells, and three imaging fields were acquired per well. For calcium imaging experiments, cells were treated with the intermediate concentration of each molecule and compared with PBS-treated controls.

### Immunocytochemistry

Cortical neurons were fixed for 10 min at room temperature in 4% PFA supplemented with 4% sucrose and washed in PBS. Primary antibodies were diluted in PBS containing 2% BSA and 0.1% Triton X-100 and applied overnight at 4 °C. The following primary antibodies were used: CTIP2 (Abcam, ab18465, 1:500), MAP2 (Abcam, ab5393, 1:1000), neurofilament (Abcam, ab4680, 1:500), and TBR1 (Abcam, ab31940, 1:500). After washing, species-specific secondary antibodies conjugated to Cy3, Alexa Fluor 488, or Alexa Fluor 647 (Jackson ImmunoResearch; 1:1000) were applied for 1 h at room temperature. Nuclei were counterstained with DAPI.

Images were acquired on a Leica SP8 confocal microscope. For each well, three independent fields of view were imaged, and for each field three optical sections were acquired as z-stacks with a step size of 5 μm. Identical imaging settings were used for all experimental conditions. MAP2- and neurofilament-positive area was quantified in ImageJ after background subtraction and application of identical thresholds across all conditions.

### Calcium Imaging

For functional calcium imaging, cortical neurons were plated at 100,000 cells cm⁻² in poly-L-ornithine- and laminin-coated 8-well ibidi chambers and analyzed after 6 weeks of terminal differentiation. Cells were loaded with the fluorescent calcium indicator Cal-520 at a final concentration of 1 μM. For each experimental condition, three independent wells were treated with the intermediate concentration of the respective molecule and three additional wells received PBS as vehicle control.

Live-cell imaging was performed at 37 °C using a Leica DMI6000B inverted epifluorescence microscope equipped with a Lumencor Aura light engine, a Leica 40× oil-immersion objective (NA 1.3), and a Hamamatsu ORCA-Flash2.8 C11440-10C CMOS camera. Images were recorded using Metafluor 7.1 software. For each well, baseline activity was recorded before treatment, and calcium activity was recorded again 1 min and 1 h after treatment. Time-lapse images were acquired every 300 ms with an exposure time of 30 ms for 10 min per recording. The same field of view was imaged at all time points. Recordings were analyzed in ImageJ, and calcium transients were quantified for event frequency, peak amplitude, and peak-to-peak interval.

### Calcium imaging data analysis

Following imaging acquisition, calcium imaging data were analyzed using Suite2p^40^ to perform cell segmentation and signal extraction. Calcium dynamics were subsequently analyzed using custom Python scripts. Raw fluorescence traces were baseline-corrected using Zhang fit (airPLS)^41^ algorithm, implemented in BaselineRemoval package^42^. Baseline-corrected fluorescence traces were then z-scored for each cell independently, according to:

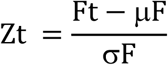

Where F_t_ is the baseline-corrected fluorescence value at frame t, μ_F_ is the mean of the baseline-corrected trace across all frames and σ_F_ the standard deviation of the baseline-corrected trace across all frames. Functional connectivity was assessed via pairwise Pearson correlation between z-scored traces and connectivity maps created using the detected cell centroids extracted from Suite2p^40^.

### Microglia Analysis

Microglial morphology was assessed by fractal analysis using a modified protocol based on previously published methods. Confocal z-stacks were converted to maximum-intensity projections, and individual microglial cells were manually isolated in ImageJ using the polygon selection tool. Only cells entirely contained within the z-stack were included. Images were thresholded, binarized, and resized to 600 × 600 pixels while preserving scale. Debris surrounding the cells was manually removed. For fractal analysis, binary images were converted to outlines and analyzed using the FracLac plugin in ImageJ. Cell area was determined from filled or outlined binary images and converted to μm² using a pixel area of 0.208 μm². Circularity was calculated as 4π × area/perimeter²

### Neuron Counting

Neuronal density was assessed in NeuN-stained sections. ROIs were defined on the basis of LnD signal, and NeuN-positive neurons within each ROI were manually counted using the ImageJ cell counter plugin. Counts were normalized to ROI area. Thirty ROIs were analyzed per animal, including 10 clot-containing ROIs, 10 clot-free ROIs in the ipsilateral hemisphere, and 10 ROIs in the contralateral hemisphere. Five animals were analyzed per group. Results were normalized to the contralateral side.

### Statistical Analysis

Data are presented as mean ± s.d. unless otherwise indicated. Statistical analyses were performed using GraphPad Prism version 8.2.1 (GraphPad Software, San Diego, CA, USA). Normality was assessed using the Kolmogorov–Smirnov test. For comparisons between two groups, Student’s t-test was used for normally distributed data and the Mann–Whitney test for non-normally distributed data. Time-course or multi-group measurements were analyzed using one-way or two-way repeated-measures ANOVA followed by Tukey’s or Sidak’s multiple-comparisons post hoc tests, as appropriate. Correlation analyses were performed using Pearson correlation. P < 0.05 was considered statistically significant.

### Transcriptomic experiments

#### Mouse single Cell RNAseq and anlaysis

Isolated cells were fixed for single cell RNA sequencing according to the manufacturer’s guidelines for whole cells (Evercode Cell Fixation V3, Parse Biosciences). Libraries were prepared using split-pool barcoding defined by the Evercode WT V3 kit (Parse Biosciences). Pooled libraries were then sequenced on Illumina NextSeq 2000 at InnovaSEQ – Cell sorting and NGS @ Caen Normandie core facilty, with a minimum of 20K reads per cell. Raw data were demultiplexed and reads were aligned to the Mus musculus reference genome (RefSeq GRCm39) using the Parse Biosciences bioinformatics pipeline module v1.6.3 in TrailmakerTM (Parse Biosciences). The dataset was processed, explored and visualized using TrailmakerTM (https://app.trailmaker.parsebiosciences.com/ Parse Biosciences, analysis completed on 2026-01-23). Unfiltered count matrices were uploaded to Trailmaker, and background was removed by setting a minimum transcripts per cell threshold on a per sample basis (threshold range: 200 to 2000). Dead or dying cells were removed by filtering barcodes with high mitochondrial content (10% cut-off). Outliers in the distribution of number of genes vs number of transcripts were removed by fitting a spline regression model (p-value 0.001). Cells with a high probability of being doublets were filtered out using the scDblFinder method. Data normalization, principal-component analysis (PCA) and data integration were then performed using Harmony. Clusters were identified using the Leiden method, and a Uniform Manifold Approximation and Projection (UMAP) embedding was calculated to visualize the results. Following quality control and preprocessing, endothelial cells were clustered based on transcriptional profiles and annotated into vascular subtypes according to established marker genes. Arteriolar endothelial cells were defined by expression of Bmx, Efnb2, Vegfc, and Sema3g with low Nr2f2 expression; venular endothelial cells by expression of Nr2f2 and Slc38a5; and capillary endothelial cells by expression of Mfsd2a and Tfrc with low Nr2f1 expression, based on previously described vascular zonation markers. An activated endothelial population was defined by increased expression of Icam1 and Vcam1. Injury-associated endothelial cells comprised the combined population of activated arteriolar, activated venular, and activated capillary endothelial cells from the ipsilateral hemisphere, whereas healthy vessel endothelial cells comprised arteriolar, venular, and capillary endothelial cells from the contralateral hemisphere. Comparative analyses were then performed between injury-associated endothelial cells and healthy vessel endothelial cells to identify transcriptional changes. Gene expression matrices were log-normalized, and missing values were treated as zero expression.

### Differential gene expression analysis

Differential gene expression analyses between traumatic brain injury and control conditions were performed using Seurat’s FindMarkers function with the Wilcoxon rank-sum test. Marker genes were identified for specific clusters and subpopulations, including endothelial cell populations isolated from cluster-specific subsets of the dataset. For cluster-level analyses, subsets of cells were extracted using Seurat’s subset function and differential gene expression was calculated between conditions within the selected cluster populations. Genes were considered significantly differentially expressed based on adjusted P values following multiple-testing correction. Differentially expressed genes (DEGs) between the injured and control cell sets were identified using the TrailMaker platform. Significant DEGs were selected using an adjusted p-value threshold of < 0.05 and subsequently subjected to Kyoto Encyclopedia of Genes and Genomes (KEGG) pathway enrichment analysis using the Enrichr web-based gene-list enrichment analysis tool to identify significantly enriched pathways.

#### Open source single-nucleus RNA sequencing data processing and quality control

Publicly available single nucleus RNA sequencing (snRNA-seq) data from human traumatic brain injury were obtained from the dataset published by Garza et al.^24^ The dataset comprises human brain tissue from patients with traumatic brain injury who underwent decompressive craniectomy and partial frontal or temporal lobe resection, as well as post-mortem tissue from non-neurological control donors, as described in the original study. Raw count matrices were imported into R (version 4.5.2) and analyzed using the Seurat package (version 5.4.0). Individual samples were loaded as Seurat objects using the CreateSeuratObject function with thresholds of min.cells = 3 and min.features = 1000 to remove genes detected in fewer than three nuclei and low-quality nuclei with fewer than 1000 detected genes. The stringent feature threshold was retained from the original study because of the high degree of damaged tissue in the TBI samples, prioritizing the inclusion of high-quality nuclei and minimizing potential bias arising from differences in tissue quality between TBI and control samples.

Quality control filtering was performed using standard Seurat workflows. The proportion of mitochondrial transcripts was calculated using the PercentageFeatureSet function. Nuclei were retained if they met the following criteria: nFeature_RNA > 1000, nCount_RNA below 100,000, and mitochondrial transcript fraction < 5%. After quality control filtering, the dataset contained 5,638 nuclei from traumatic brain injury samples (n = 12 patients; average detected genes per nucleus = 2,588) and 7,144 nuclei from control samples (n = 5 individuals; average detected genes per nucleus = 3,574).

Gene expression counts were normalized using Seurat’s NormalizeData function and highly variable genes were identified using FindVariableFeatures with 3000 variable features. Dimensionality reduction was performed using principal component analysis (RunPCA) using the variable genes and calculating 30 principal components.

#### Human scRNA sequencing analysis

The open-source data were re-analyzed from GEO (GSE209552)^24^ using Trailmaker (Parse Biosciences). Processed count matrices (barcodes, features and matrix files) were imported as individual samples and analyzed using the standard Trailmaker pipeline. Cells were subjected to quality-control filtering and doublet detection using Trailmaker default procedures, and normalized expression values were generated for downstream analyses.

Unsupervised dimensionality reduction and Leiden clustering were performed in Trailmaker, and clusters were annotated based on canonical marker gene expression. Brain endothelial cell populations were identified by enrichment of PECAM1, VWF, KDR/FLT1, ENG and CLDN5 and manually curated to minimize contamination from mural (PDGFRB, RGS5), fibroblast (COL1A1, LUM) and immune cell populations. Endothelial cells were further clustered into BEC subpopulations, and subsequent analyses were restricted to the BEC2 cluster.

Normalized expression values for BEC2 cells were exported from Trailmaker as a gene-by-cell expression matrix and used for downstream endothelial functional scoring. All genes included in the functional gene sets were confirmed to be detected in the normalized expression matrix and were retained irrespective of dispersion thresholds to preserve low-abundance but biologically relevant endothelial transcripts. For each endothelial cell, composite gene-expression scores were calculated as the unweighted mean normalized expression of predefined gene sets: pro-coagulatory (VWF, EDN1, ANGPT2, ENG), thrombolytic (PLAT, ANXA2, ANXA11, SERPINE1), barrier/junctional (CLDN5, PECAM1, TJP1, OCLN) and transcytotic (CAV1, CAV2, PLVAP, CAVIN1). An integrated endothelial bias score was calculated for each cell as the transcytosis score minus the barrier/junctional score. Comparisons between control and TBI endothelial cells were performed using two-tailed Mann–Whitney tests in GraphPad Prism. Data were visualized using violin plots with individual endothelial cells overlaid as data points.

### Data integration, clustering, and visualization

To account for batch effects across samples, datasets were integrated using the Harmony algorithm. Dimensionality reduction was performed on the Harmony-corrected embeddings and the first 15 dimensions were used for downstream analyses. A shared-nearest-neighbor graph was constructed using the FindNeighbors function with dims = 1:15, and cell clusters were identified using the Louvain algorithm implemented in FindClusters with a resolution parameter of 0.3.

Two-dimensional visualization of cellular populations was generated using Uniform Manifold Approximation and Projection (UMAP) with the RunUMAP function applied to the Harmony embeddings using dims = 1:15.

Cell types were annotated based on the expression of canonical marker genes from previously published human brain single-cell datasets. Endothelial cells were identified based on expression of vascular markers including PECAM1 (CD31), CLDN5, and VWF. After subsetting endothelial populations, a total of N = 89 endothelial nuclei were retained for downstream analyses.

### Human TBI cohort and clinical data

A retrospective clinical cohort study was performed at the Department of Neurosurgery, Ludwig-Maximilians-University (LMU) Munich. Patients aged >65 years who were admitted with a diagnosis of traumatic brain injury (TBI) between 2010 and 2016 were screened for inclusion. A total of 550 patients were initially identified. After exclusion of patients with secondary traumatic injuries (n = 190) and those admitted more than 6 h after injury (n = 131), 229 patients with isolated acute TBI remained for analysis. Within this cohort, 114 patients were receiving anticoagulant or antiplatelet medication at the time of injury, whereas 115 patients had no anticoagulant treatment. Clinical data including demographics, comorbidities, injury characteristics, radiological findings, laboratory parameters, treatment variables, and clinical outcomes were extracted from electronic medical records. (Extended Data Fig. 7 b,c). Clinical characteristics, neuroimaging findings and longitudinal laboratory measurements were extracted from the available clinical records.

### Ethics statement

This study was approved by the institutional Ethical Committee of Munich University Medical Faculty (protocol # 21-0712) and conducted in accordance with the Declaration of Helsinki.

### Clinical Data Collection

For each patient, demographic variables (age and sex), injury severity parameters, and clinical outcomes were recorded. TBI severity was classified based on the Glasgow Coma Scale (GCS) at admission as mild (GCS 13–15), moderate (GCS 9–12), or severe (GCS 3–8). Comorbidities including arterial hypertension, coronary heart disease, atrial fibrillation, chronic obstructive pulmonary disease, liver disease, kidney failure, dementia, and prior stroke or transient ischemic attack were documented. Radiological characteristics were assessed from admission computed tomography scans and included the presence of contusions or subarachnoid hemorrhage, epidural hematoma, subdural hematoma, midline shift, and radiological clot thickness score (RCTS). Injury severity score (ISS) and clinical course variables including intensive care unit stay, hospital stay duration, surgical intervention, and mortality were also recorded.

### Longitudinal analysis of haemostatic parameters

Routine coagulation and hematological parameters were obtained from clinical laboratory measurements during hospitalization. These included hemoglobin levels, platelet counts, prothrombin time (Quick), activated partial thromboplastin time (aPTT), fibrinogen concentration, antithrombin levels, and D-dimer levels. For longitudinal analyses, laboratory values were evaluated at predefined clinical time points. To characterize the intrinsic haemostatic response to acute TBI while minimizing the influence of pre-injury antithrombotic medication, patients were stratified according to pre-injury use of direct oral anticoagulants (DOACs) and/or antiplatelet agents (Extended Data Fig. 7b). Longitudinal trajectories of aPTT, Quick, platelet count, fibrinogen and antithrombin were examined in patients not receiving either DOAC or antiplatelet therapy. Measurements were grouped according to predefined post-injury sampling intervals. For visualization of the relative temporal trajectories in Fig. 5a, each parameter was expressed relative to its corresponding healthy reference value, defined as 100%. Additional longitudinal laboratory parameters, including D-dimer and haemoglobin, were analysed in the broader clinical cohort (Extended Data Fig. 7d). To assess whether pre-injury antithrombotic treatment modified the acute fibrinogen response, fibrinogen concentrations were additionally compared between patients receiving DOAC and/or antiplatelet therapy and patients receiving neither treatment across the corresponding post-injury sampling intervals. Data are presented as median with interquartile range (IQR), and between-group comparisons at individual time points were performed using two-sided Mann–Whitney *U* tests.

### Definition of early post-TBI coagulopathy

For the exploratory translational analysis, early coagulopathy was defined *a priori* as INR >1.2 and/or aPTT >35 s in any available blood sample obtained 1–11 h after admission. Patients with available coagulation measurements during this interval who did not exceed either threshold were classified as non-coagulopathic. Patients without sufficient measurements for classification within the predefined interval were excluded from the coagulopathy-stratified analysis. Of the 229 patients in the clinical cohort, 222 were evaluable according to these criteria, comprising 63 patients with and 159 patients without early coagulopathy. Continuous clinical variables were compared between coagulopathy groups using Mann–Whitney U, and categorical variables using χ²/Fisher. Continuous variables are reported as median (IQR) and categorical variables as *n/N* (%). Statistical significance was defined as *P* < 0.05.

### Molecular imaging probes

To evaluate the endothelial expression of VCAM-1 within the cerebral vasculature, microparticles of iron oxide (MPIO) conjugated with an anti-VCAM-1 antibody were used. MPIO synthesis, antibody coupling, and intravenous dosing configurations were performed exactly as previously described by Gauberti et al.^43^

### In vivo MRI and physiological monitoring

A total of 2 mice were used. MRI was performed using a 7-Tesla preclinical scanner (BioSpec 70/16; Bruker Biospin, Ettlingen, Germany) running ParaVision 6.0.1 software, equipped with a quadrature transmit/receive surface coil for mouse brain imaging (RF COIL 300 1H M.BR QSN, Model T11204V3; Bruker BioSpin). To track the spatiotemporal evolution of the lesion and the inflammatory response via VCAM-1 expression, animals underwent longitudinal MRI sessions at two distinct time points: early acute (+6 hours post-TBI) and delayed (+24 hours post-TBI).

Throughout each imaging session, mice were anesthetized using isoflurane (1.5%–2% in 100% oxygen) adjusted dynamically to maintain a stable respiratory rate of approximately 90 breaths per minute. Continuous physiological monitoring was performed to ensure stable core body temperature and respiratory rate during the entire acquisition.

#### MRI sequence parameters

For lesion characterization, axial T2-weighted images were acquired using a multi-slice multi-echo spin-echo sequence. The axial slice orientation was selected to ensure a homogeneous signal profile across each field of view, thereby mitigating the depth-dependent signal gradient inherent to surface coil reception profiles. The sequence parameters were as follows: repetition time (TR) = 3500 ms, echo time (TE) = 40 ms, number of echoes = 8, flip angle = 90°, number of averages = 2, acquisition matrix = 184 × 256 (reconstructed to 256 × 256), field of view (FOV) = 17.9 × 17.9 mm², in-plane resolution = 70 × 70 µm², 18 contiguous slices, slice thickness = 0.5 mm, and slice separation = 0.5 mm.

For MPIO detection, high-resolution axial T2*-weighted gradient-echo images (FLASH) were performed immediately before and after MPIO injection. As with the T2w sequences, an axial orientation was chosen to avoid within-slice signal intensity gradients caused by the dorsal-to-ventral sensitivity drop-off of the surface coil, ensuring reliable quantification of MPIO-induced signal voids. To minimize confounding susceptibility artifacts arising from deoxyhemoglobin within the cerebral vasculature, acquisitions were conducted under 100% oxygen ventilation^44^. The sequence parameters were as follows: TR = 100 ms, TE = 3 ms, number of echoes = 1, flip angle = 30°, number of averages = 1, acquisition matrix = 128 × 128 (reconstructed to 256 × 256), FOV = 25 × 25 mm², in-plane resolution = 98 × 98 µm², 5 slices, and slice thickness = 1.0 mm.

#### Image processing and signal void quantification

Image processing and quantification of MPIO-induced hyposignals were performed using ImageJ/Fiji software. First, image contrast was enhanced by 0.35% with intensity normalization applied across all slices to standardize baseline gray levels. To reduce high-frequency voxel noise and smooth the susceptibility blooming effect along the slice axis, a 3D Minimum Filter was applied with a neighborhood radius configuration of X = 0, Y = 0, and Z = 1 pixel (effectively processing adjacent slice vectors).

Automated segmentation of the hypointense areas was achieved using an Otsu thresholding, restricting the selected normalized intensity range between 0.007 and 0.337 to accurately isolate MPIO clusters from brain tissue. Intracerebral signal void areas were then measured within manual regions of interest delineating the ipsilateral and contralateral hemispheres. To isolate the specific contrast generated by the microparticles from pre-existing tissue damage, hemorrhages, or baseline anatomical artifacts, a differential correction formula was applied to each hemisphere:

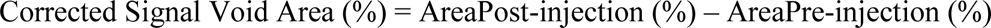

### Statistical Analysis

Statistical analyses were performed using GraphPad Prism (GraphPad Software). Continuous variables are presented as mean ± standard deviation unless otherwise indicated. Comparisons between patient groups (patients with versus without anticoagulant medication) were performed using unpaired two-tailed t-tests for normally distributed variables. For longitudinal laboratory measurements, mixed-effects models using restricted maximum likelihood (REML) estimation were applied to account for repeated measurements and missing values. Time point was treated as a fixed effect, and patient identity was included as a random effect. Sphericity was not assumed and the Geisser–Greenhouse correction was applied when appropriate. When significant effects were detected, Dunnett’s multiple-comparisons test was used to compare post-injury time points with baseline values. Categorical variables were compared using χ² tests. Statistical significance was defined as P < 0.05.

## Acknowledgements

The authors thank Georg Kislinger for tissue preparation in CLEM experiment. Single cell RNAseq experiments were performed at InnovaSEQ – Cell sorting and NGS @ Caen-Normandie Facility. The European project EquipInnovCaen2022-RNAseq is funded by the European Union within the framework of the Operationnal Programme ERDF/ESF 2014-2020.The European project SLIM is funded by the European Union within the framework of the Operationnal Programme ERDF/ESF 2021-2027. This work was supported by DFG 457586042, ANR-DFG MICROSTROKE, MSCP program LMU.

---

**Extended data Figure 1.**
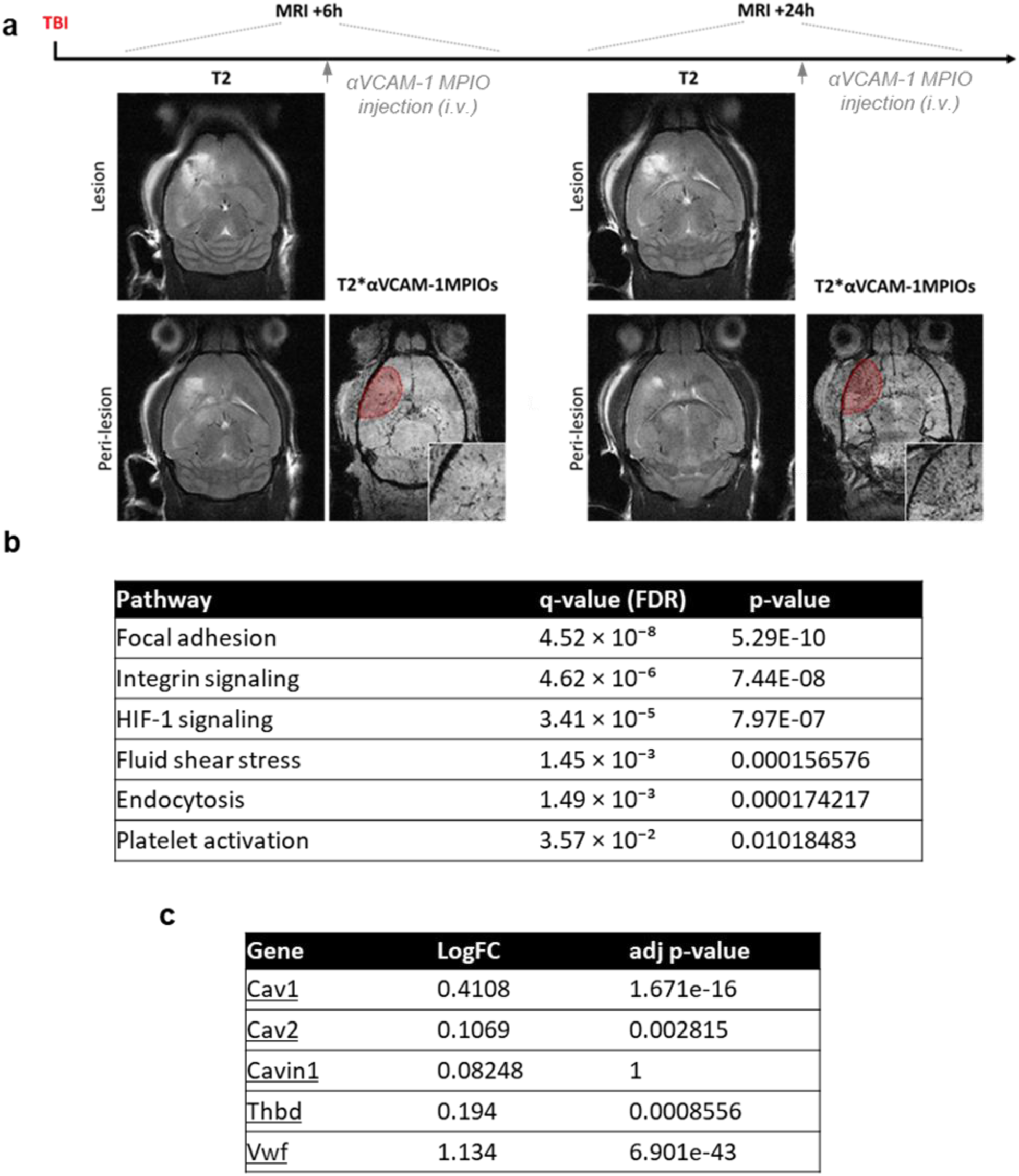
Molecular MRI and scRNA seq test. **a,** Experimental timeline and representative axial T2*-weighted images before (Pre-MPIO) and after (Post-MPIO) αVCAM-1 MPIO injection at 6 and 24 hours on the same animal (n=2). The structural T2-weighted image outlines the lesion sites, whereas the corresponding T2*-weighted image highlights acute development of perilesional VCAM-1 expression (red area). **b,** q-values (FDR (False Discovery Rate)) and p-values of KEGG pathways from injured BEC at figure 1 g. **c,** log2FC and adjusted p-values for key genes (figure 1 h-i). Analysis was performed using per-cell Wilcoxon test.

**Extended data. Figure 2.**
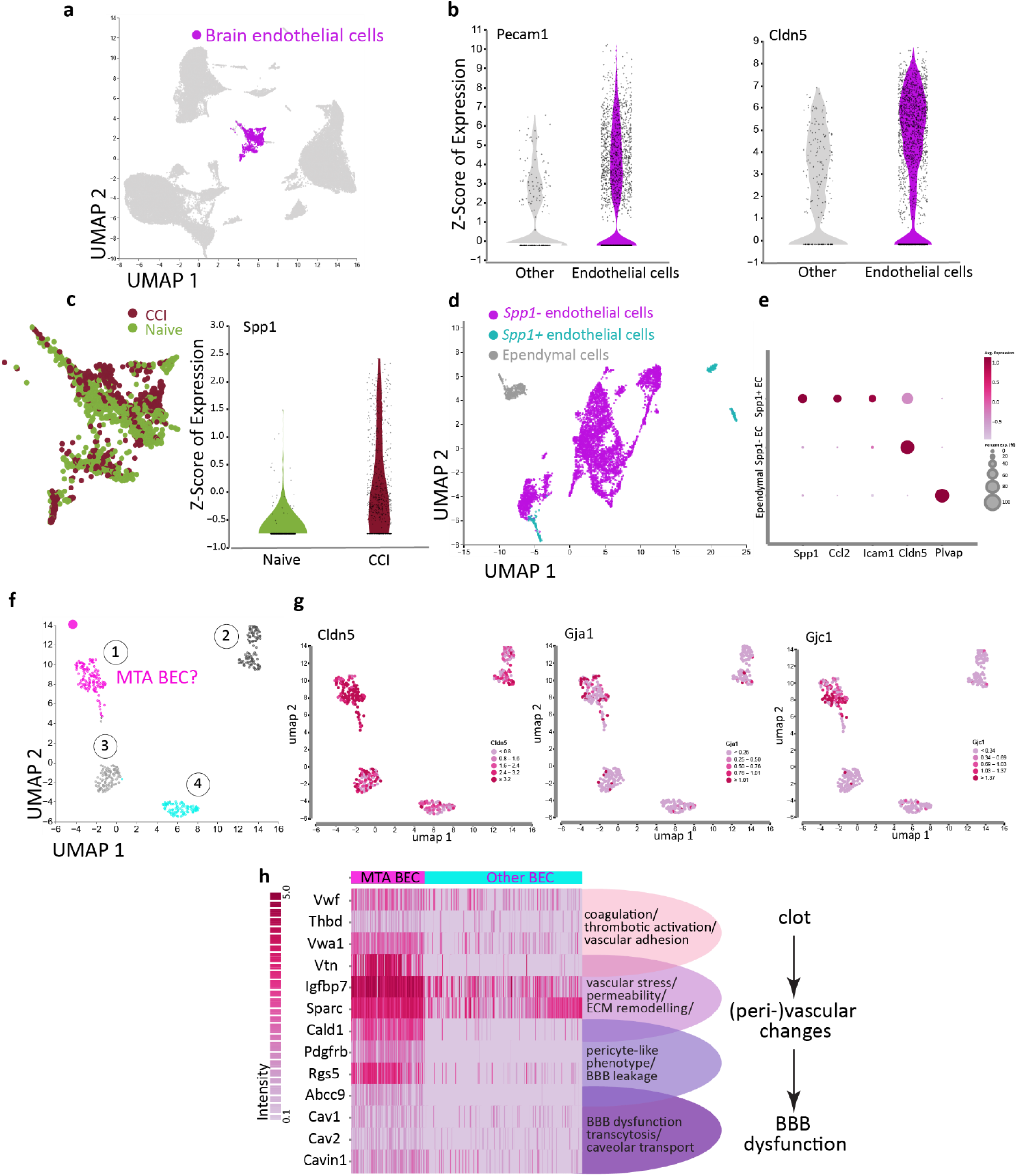
Identification and characterization of a transcriptionally distinct injury-related Spp1⁺ endothelial cell population 24 h following controlled cortical impact (CCI). **a,** UMAP plot highlighting brain endothelial cells (magenta) among all brain cell types in mice of two groups (Naïve and CCI) processed from original dataset Jha, Ruchira M. et al. **b,** Violin plots showing elevated expression endothelial markers. **c,** Endothelial cells cluster of two conditions (Naïve and CCI) and increased Spp1 expression in CCI vs. naive conditions (right). **d,** UMAP of further processed endothelial cells of CCI group reveals Spp1⁺ endothelial cells (cyan) and Spp1⁻ endothelial cells (magenta), showing additionally a transcriptional separation of ependymal cells (gray). **e,** Dot plot showing that Spp1⁺ endothelial cells are enriched by expression of Icam and inflammatory genes, suggestive a pool of injured BEC, while Spp1^-^ are not. **f,** UMAP clustering of injured BEC identifies a distinct cluster (1) with upregulation of microthrombus-associated (MTA) genes. **g,** UMAP feature plots showing the expression of Cldn5, Gja1, and Gjc1 within the Spp1⁺ endothelial subcluster identified 24 h after CCI. The co-expression of canonical endothelial marker Cldn5 confirms vascular identity, whereas selective enrichment of the gap-junction genes Gja1 and Gjc1 define post-capillary venules, suggesting that these injury-associated endothelial cells originate from the venular compartment, the region most prone to microthrombus formation and BBB leakage. **h,** Heatmap showing elevated expression of genes involved in microthrombus formation, vascular stress, pericyte-like transition and caveolar transcytosis reflecting a possible progression from microthrombus to BBB breakdown. Original bulk RNA-seq data are from Jha, Ruchira M. et al.

**Extended data. Figure 3.**
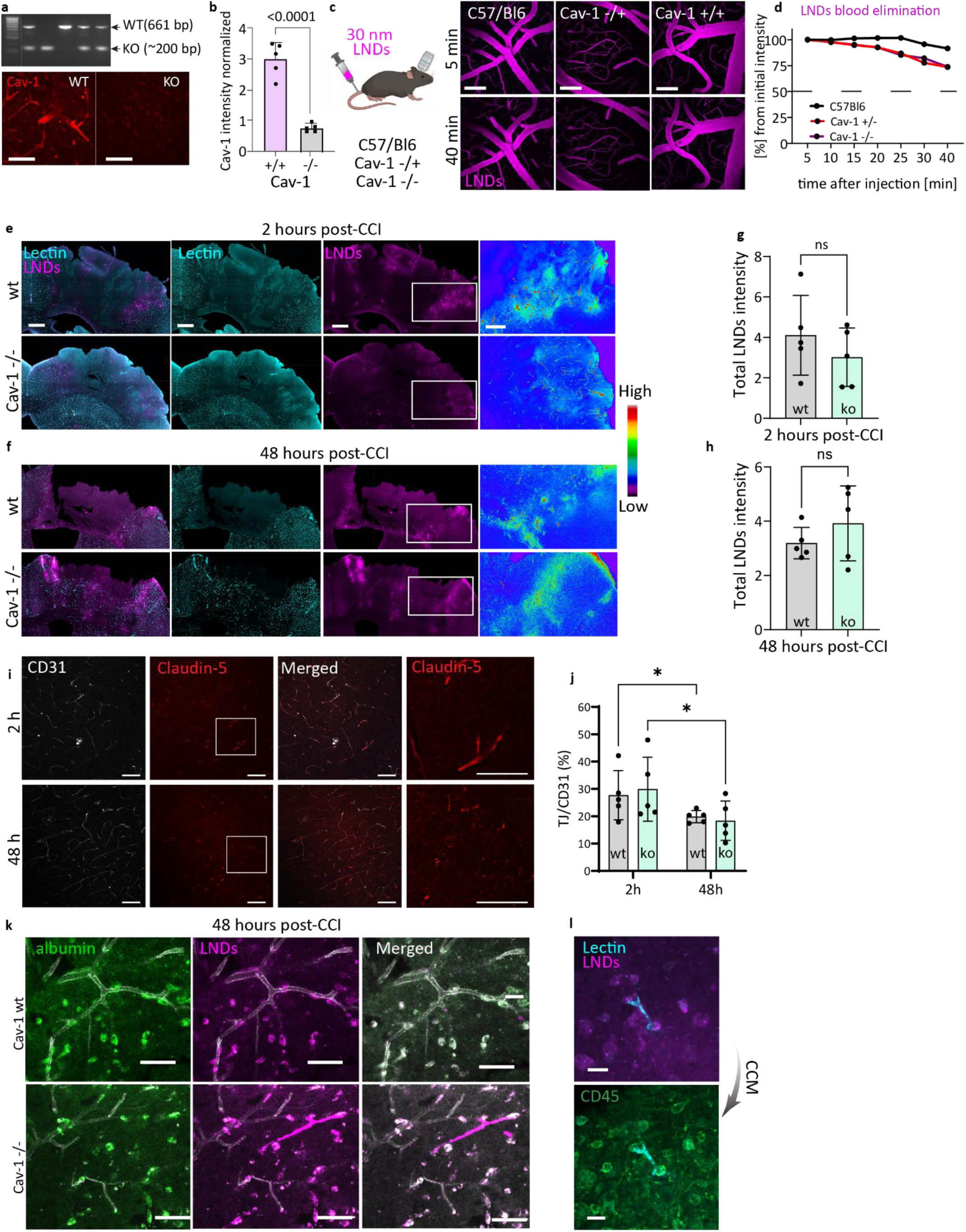
Cav-1 related dynamics in MTi and BBB dysfunction. **a-b,** PCR-based genotyping confirming Cav-1 knockout in homozygous (Cav-1 ko/ko) mice compared to wild-type (Cav-1 wt/wt) controls, and validation of Cav-1 deletion by immunohistochemistry showing reduced Cav-1 signal in brain vasculature of Cav-1 ko/ko mice and its quantification (b). Scale bar: 50 µm. Data are Mean ± SD; n = 5 mice, unpaired t-test. **c-d,** Representative time-lapse two-photon microscopy through a glass cranial window showing circulation of 30 nm LNDs in cortical vasculature after systemic injection in C57Bl/6N, Cav-1 wt/ko, and Cav-1 ko/ko mice. Scale bar: 100 µm. **d,** Quantification of LND circulation kinetics over 40 minutes across genotypes (mean of 3 ROI; n = 1). **e-f,** Representative confocal tile images of the ipsilateral cortex 2 h (e) and 48h (f) post-CCI showing gross extravasation and accumulation of LNDs in Cav-1 ko and Cav-1 wt mice. Scale bar: 500 µm; zoomed 200 µm. **g-h,** Quantification of extravasation intensity 2 h (g) and 48 h (h) post-CCI across the genotypes (Mean ± SD; n = 5 mice per group; unpaired t-test). **i-j,** Representative confocal images of tight junctions (claudin-5) and their quantification 2 and 48 h after CCI across the genotypes. Scale bar: 50 µm. (Mean ± SD; n = 5 mice per group; two-way ANOVA, followed by Šídák’s multiple comparisons test.). **k,** Local extravasation pattern 48h post-CCI across genotypes. Scale bar: 50 µm. **l,** Representative of CCM showing intracellular localization of extravasated LNDs within CD45⁺ immune cells. Scale bar: 20 µm.

**Extended data Figure 4.**
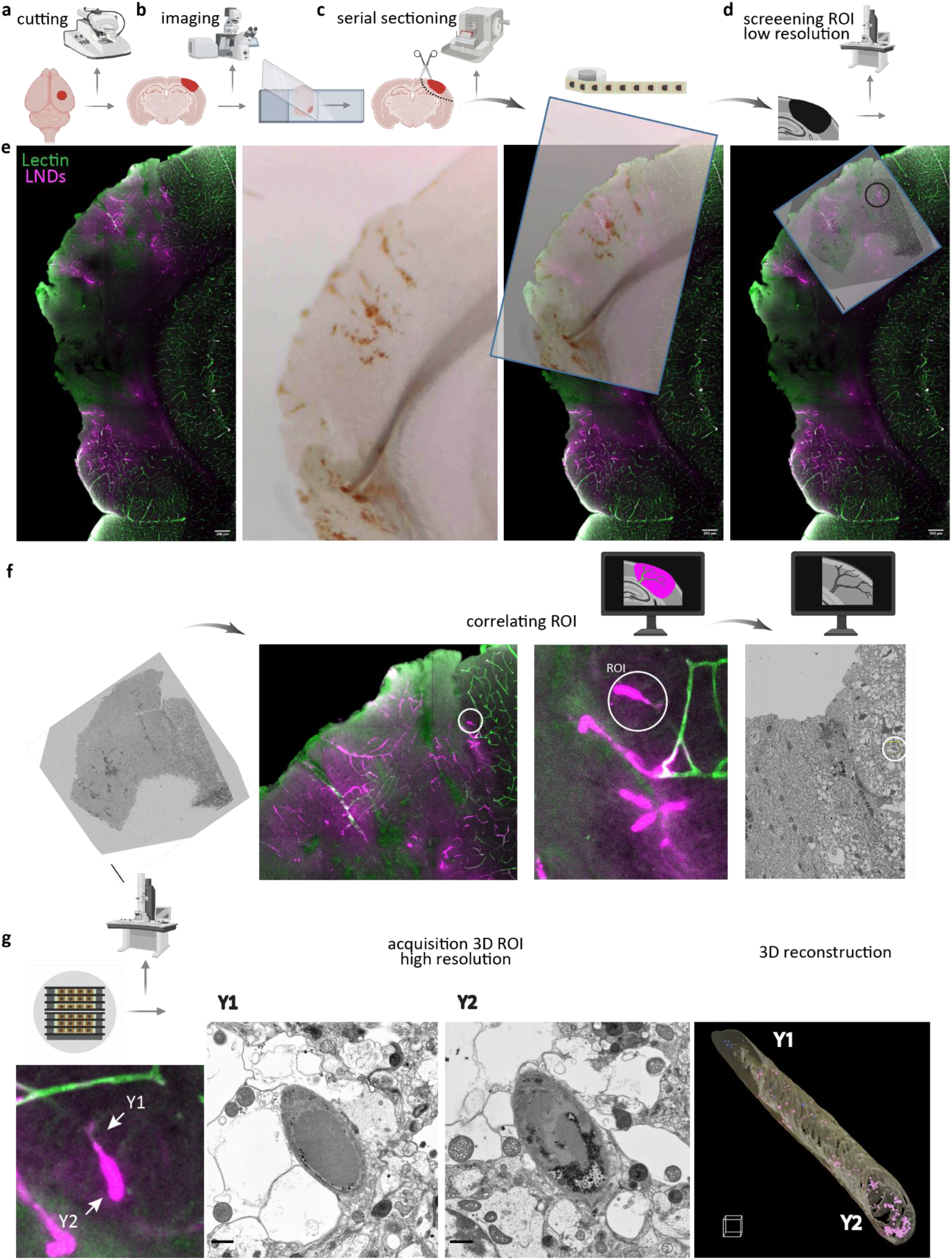
Targeted CLEM pipeline for ultrastructural analysis of microthrombi and LND extravasation after CCI. **a,** Schematic of the targeted CLEM workflow following experiment presented at Fig. 3d. After fixation with 2.5% glutaraldehyde of injured mouse brain, coronal vibratome sections are generated and placed onto glass slides with a coverslip gently laid on top for confocal imaging of LNDs and lectin-labelled vessels. **c,** The section is then recovered by immersion into a petri dish with PBS, and the previously imaged cortical region of interest (ROI) is embedded into resin and dissected from the vibratome slice using Serial ultramicrotomy with tape collection (ATUM)**. d,** The consecutive screening is performed of the section mounted on silicon wafers for subsequent targeted high resolution scanning electron microscopy (SEM). **e-f,** Correlative mapping between confocal and SEM datasets allows precise re-identification of the originally imaged LND-positive structures. **g,** Higher-magnification confocal images of two selected LND-labelled structures (Y1 and Y2, arrows) and corresponding serial SEM images as well as 3D reconstruction (magenta – intraluminal NPs; blue – intracellular). Scale bars: confocal, 200 μm (overview), 10 μm (zoom); SEM panels, 1 μm, 3D voxel - 2 μm.

**Extended Data Figure 5.**
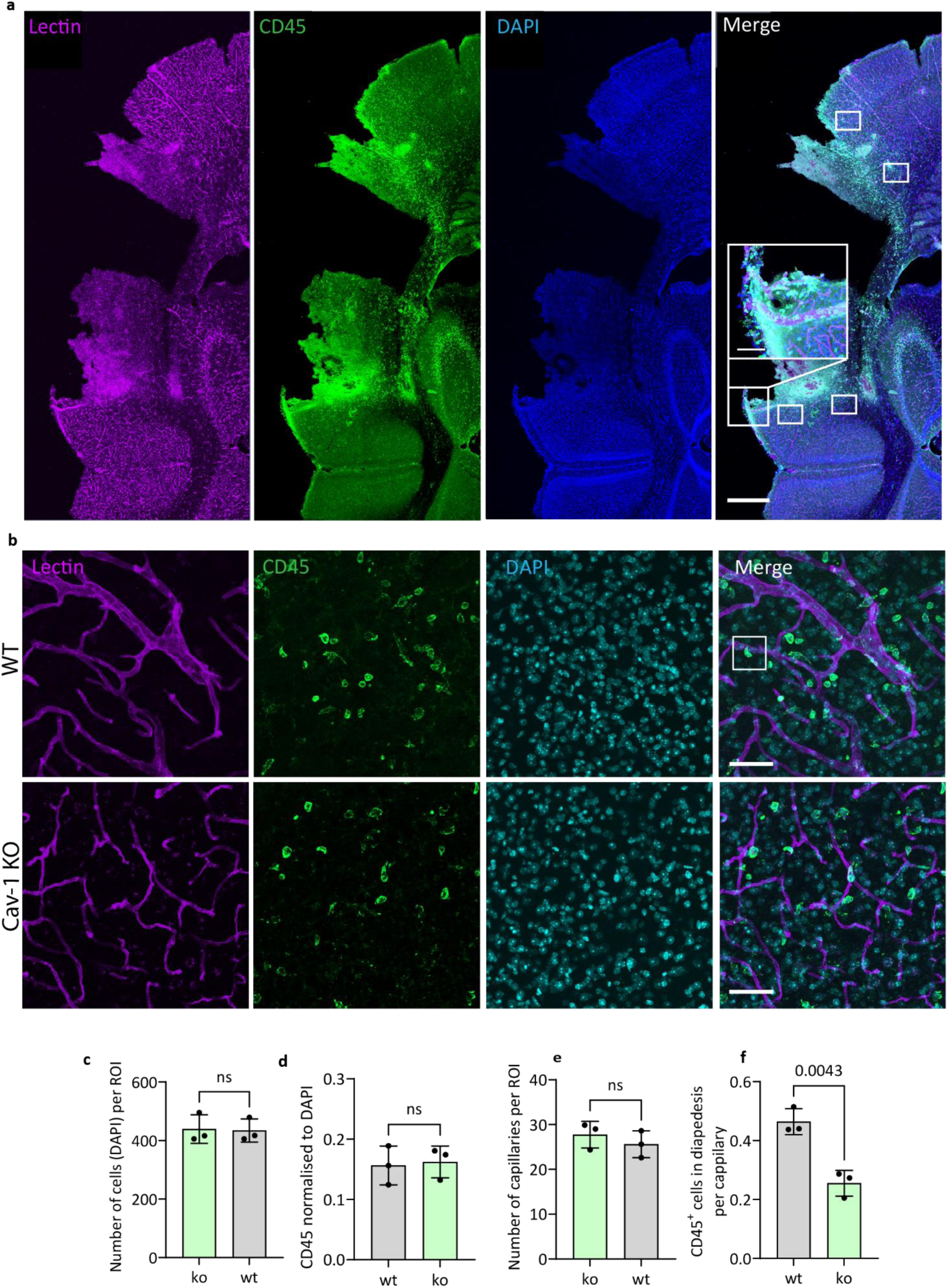
Immune cell trafficking in the perilesional cortex 48 h after CCI. **a,** Representative confocal images of CD45⁺ immune cells in the cortical lesion and perilesional area in WT animals. **b,** Upper panel - higher-magnification region of interest (ROI) from the perilesional area showing a CD45⁺ cell undergoing diapedesis (white box); lower panel - homotopical ROI from Cav-1 KO mice. **c,** Quantification of total cell number (DAPI⁺ nuclei) in the perilesional area showing no difference between genotypes. **d,** Fraction of CD45⁺ immune cells among total DAPI⁺ cells in the perilesional area showing no difference between genotypes. **e,** Quantification of the number of capillaries within ROIs showing no difference between genotypes. **f,** Number of CD45⁺ cells undergoing diapedesis per capillary showing a reduction in Cav-1 KO mice. Data are mean ± SD; n = 4 ROIs from 3 sections per mouse, 3 mice per genotype; unpaired two-tailed t-test. Scale bars, 1 mm (a); 100 µm (insert in a); 50 µm (b).

**Extended Data Figure 6.**
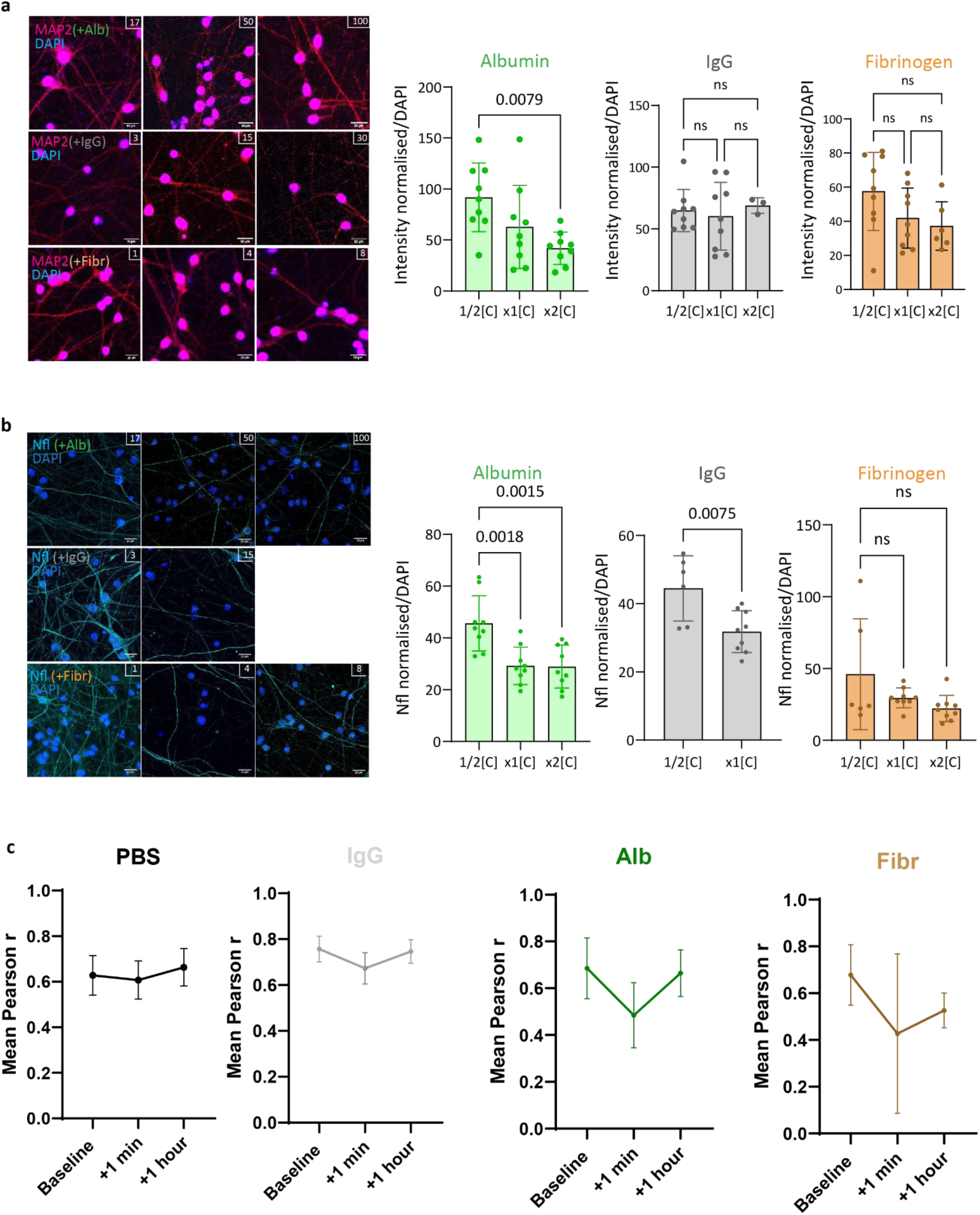
Dose-and size-dependent effects of blood-borne proteins on human iPSC-derived neurons. Representative MAP-2 **(a)** and Nfl **(b**) images and quantification of iPSC-derived neurons treated with IgG, albumin, or fibrinogen at 0.5×, 1×, and 2× human plasma-equivalent concentrations [C]. Data are mean ± SD; n=9 ROIs from 3 wells per condition; One-way ANOVA. Scale bar, 20 µm. **c,** Quantification of mean Pearson r of connectivity of iPSC-derived neurons after exposure to blood born components. Data are mean ± SD; n=3-4.

**Extended Data Figure 7.**
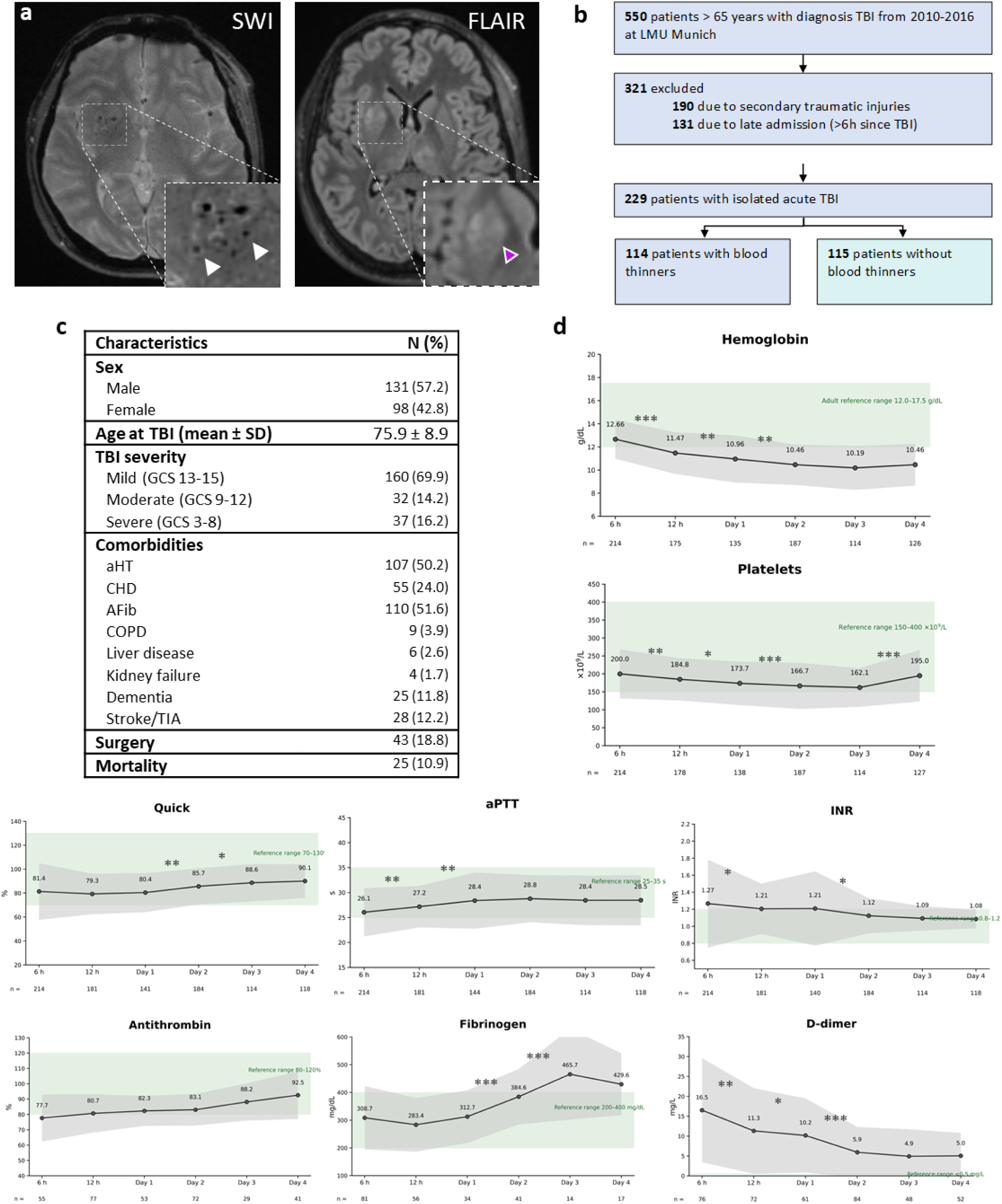
MRI and characteristics of human acute TBI cohort. **a**, Representative susceptibility-sensitive brain MRI illustrates haemorrhagic lesions associated with TBI (white arrows) and correspondent FLAIR image showing tissue damage (magenta arrows). **b,** Cohort stratification steps according to type of TBI, admission time and pre-injury blood antiplatelets and anticoagulants use. Patients were divided into antiplatelet and anticoagulants takers and non-takers for comparative analyses. **c,** Characteristics of human isolated TBI cohort from LMU hospital. **d,** Longitudinal blood-based signatures of coagulation activation, consumption, and counter-regulation following acute TBI. Data are presented as mean ± SD. Green shaded areas indicate the respective clinical reference ranges. Longitudinal changes were assessed using linear mixed-effects models with time as a fixed effect and patient as a random intercept. P values were adjusted using the Holm method. D-dimer values were log-transformed before analysis. *P < 0.05, **P < 0.01, ***P < 0.001. Not significant – not shown. Numbers below the x-axis indicate the number of patients with available measurements at each time point.

**Extended Data Figure 8.**
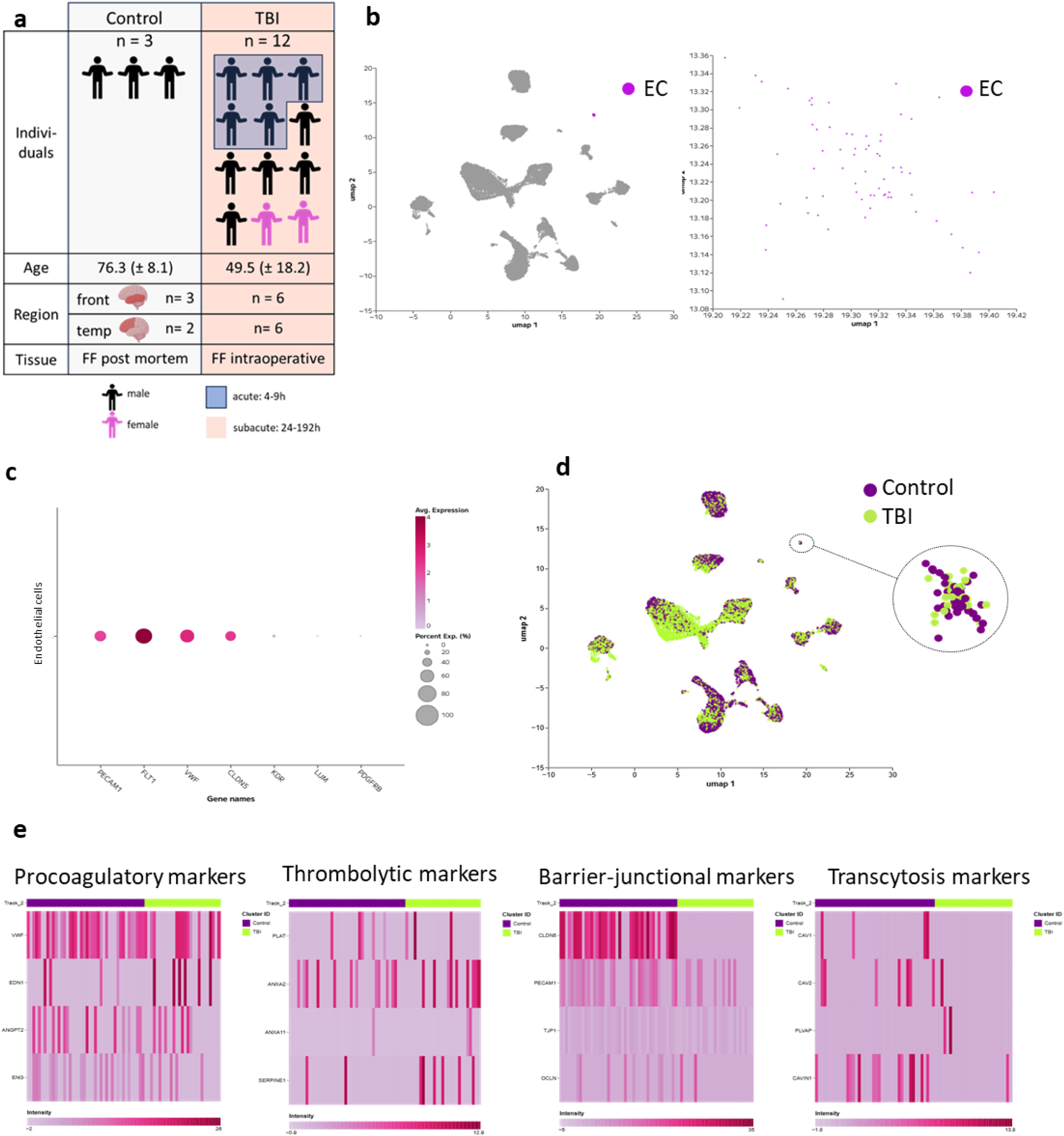
Endothelial transcriptional signatures in human traumatic brain injury. Single-nucleus RNA sequencing (snRNA-seq) data from human cortical tissue (Garza et al.)^24^ were analyzed to assess endothelial gene expression signatures in control and traumatic brain injury (TBI) samples. **a,** Description of the cohort. **b,** UMAP representation of all sequenced nuclei, with endothelial cells (ECs) highlighted, showing clustering of the endothelial population within the broader cortical cell landscape. **c,** Expression of endothelial marker genes within the identified EC cluster, confirming endothelial identity. **d,** UMAP visualization of endothelial cells separated by condition (control versus TBI), illustrating injury-associated transcriptional differences within the EC population. **e,** Heatmaps showing normalized expression of genes associated with pro-coagulatory, thrombolytic, barrier/junctional, and transcytotic endothelial signatures in ECs from control and TBI samples.

## Notes

### Competing Interest Statement

The authors have declared no competing interest.

### Summary of Updates

We added one panel to figure Extended Data Figure 5 and statistics to Extended Data Figure 7. And removed paragraph from Material and Metods which potentially could clash with future patent applilcation.

